# Spherical Code of Retinal Orientation-Selectivity Enables Decoding in Ensembled and Retinotopic Operation

**DOI:** 10.1101/2025.01.08.631850

**Authors:** Dimitrios D. Laniado, Yariv Maron, John A. Gemmer, Shai Sabbah

## Abstract

Selectivity to orientations of edges is seen at the earliest stages of visual processing, in retinal orientation-selective ganglion cells (OSGCs), thought to prefer vertical or horizontal orientation. However, as stationary edges are projected on the hemispherical retina as lines of longitude or latitude, how edge orientation is encoded, and decoded by the brain, is unknown. Here, by mapping the OS of thousands of OSGCs at known retinal locations in mice, we identified three OSGC types whose preferences match two longitudinal fields, and a fourth type matching two latitudinal fields, the members of each field-pair being non-orthogonal. A geometric decoder revealed that two OS sensors yield optimal orientation decoding when approaching the deviation from orthogonality we observed for OSGC field-pairs. Retinotopically-organized decoding generated type-specific variation in decoding efficiency across the visual field. OS tuning was greater in the dorsal retina, possibly reflecting an evolutionary adaptation to an environmental gradient of edges.

## Introduction

A key property of neurons in the visual system is a differential selectivity to edges of particular orientations. Extending throughout the visual system, orientation selectivity (OS) is required for visually-guided behaviors, object detection, and face recognition^1–17^. Discovered in the cat’s primary visual cortex (V1), OS was suggested to originate from feedforward orientation-indifferent excitatory input from the dorsal division of the retinorecipient lateral geniculate nucleus (dLGN)^18,19^. The ‘feedforward model’^20,21^, along with alternative models^22,23^, successfully explains OS created *de novo* in the cortex, either through non-OS dLGN input, or via intracortical interactions. However, existing models do not account for the observation that the mouse dLGN provides OS inputs to V1^24–26^, and harbors orientation selective neurons with a greater preference bias towards the cardinal orientations than in V1^27–30^. This, together with the independence of dLGN OS of corticothalamic feedback^27–29,31^, and thus of cortical OS, raises the question of whether retinal OS plays a role in the dLGN OS. At least some OS in the dLGN, and also in V1, might be the product of input from retinal *direction*-selective ganglion cells (DSGCs), converging on dLGN neurons^24,29,32^. However, retinal input from *orientation*-selective ganglion cells (OSGCs) may also endow the dLGN and subsequently the cortex with OS^32,33^. OS in the superior colliculus (CS), a sensorimotor hub integrating visual and other sensory information to drive reflexive behaviors, might also be grounded in OSGCs signaling^34–36^.

Among mammals, OSGCs have been best characterized in the rabbit^37^ and mouse^28,38–42^, where the orientation preferences of OSGCs were found to be biased toward either one or two of the cardinal orientations – vertical or horizontal (but see Baden et al.^42^). Importantly, according to the current paradigm, OSGCs of a given type are divisible into two populations, each responding most effectively to the “same orientation” everywhere in the retina, and, the preferred orientations of the two populations are perpendicular to one another. This widely held idea, however, has never been adequately defined in relation to the appropriate spherical geometry of the retina.

On geometrical grounds alone, any stationary real-world edge (line), or segments thereof, may be locally approximated as a line of longitude or latitude (Figure 1A). The projection of such a line on the hemispherical retina would likewise follow locally lines of longitude or latitude (Figure 1B). Spherical geometry also dictates that similar-orientation edges at other visual-space locations, will be projected on other retinal locations at orientations aligned with the local lines of longitude or latitude (Figure 1C). Deviation from such spherical logic would degrade the retinal representation of edges longer than the receptive fields of OSGCs (5.9-10.3° visual angle; see methods)^38,43^. These stationary longitudinal and latitudinal fields represent a snapshot of the optic flow patterns generated when the animal translates along (longitude), or rotates about (latitude) a given axis, recently used to explain the preferred directions of DSGCs^44,45^. We postulated that identifying a spherical geometry model that explains OSGCs preferences, will provide insight on the retinal information transmitted to V1 via the dLGN, and possibly to other OSGC-recipient regions. Additionally, determining whether the model best explaining OSGCs preferences, is similar to the model best fitting DSGCs preferences, could provide insight into the developmental mechanisms generating retinal OS and DS.

**Figure 1.**
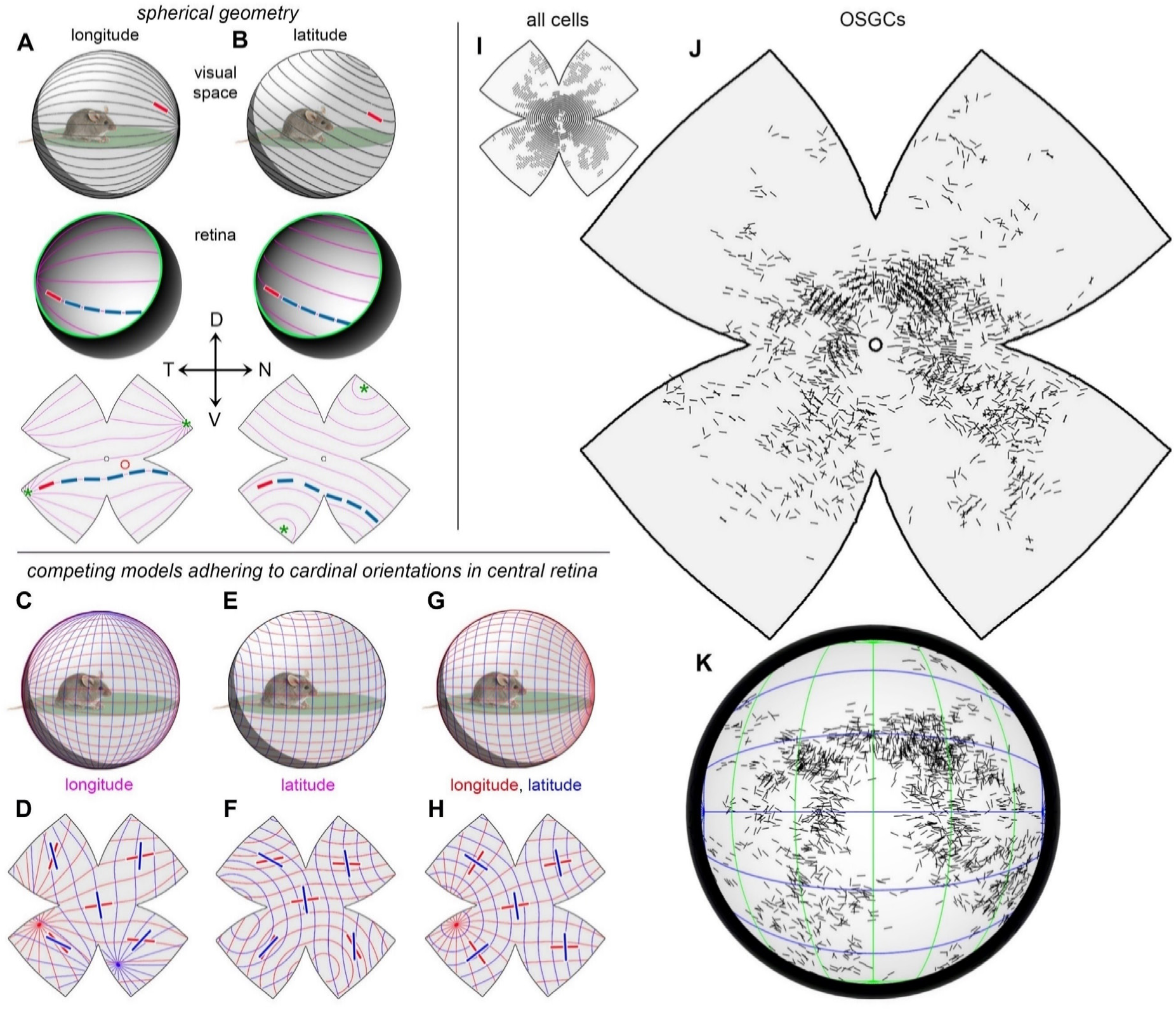
Orientation preferences of OSGCs vary across the retina A,B, top. A stationary real-world edge segment (red line) may be locally approximated as a line of longitude or latitude. **A,B, middle, bottom.** The projection of a real-world edge follows lines of longitude or latitude, on a hemispherical (**middle**) and flattened (**bottom**) retina. The projection of similar-orientation edges at adjacent visual-space locations (blue lines), alignes with local longitude or latitude lines. Asterisks, center of longitudinal (**A, bottom**) or latitudinal (**B, bottom**) fields. N, nasal; D, dorsal; T, temporal; and V, ventral. **C-H.** Three competing spherical models adhering to cardinal orientations (horizontal and vertical) in the central retina, viewed in the visual space around the animal (**C,E,G**) and projected onto the retina (**D,F,H**). Orientation-selectivity (OS) preferences of orientation-selective ganglion cells (OSGC) would follow two longitudinal fields, rotated by 90°, the fields being either both longitudinal, *i.e.*, a ‘longitudinal geometry’ model (**C,D**); both latitudinal, *i.e.*, a ‘latitudinal geometry’ model (**E,F**); or one longitudinal and one latitudinal, *i.e.*, a ‘hybrid geometry’ model (**G,H**). **I.** Locations of all imaged cells (dots) mapped onto a computationally-flattened retina. **J,K.** Topographic dependency of OSGC’s local OS preferences, displayed on a standardized flattened retinal map (**J**), and on a projected spherical retina (**K**). For reference, green and blue lines in (**K**) represent orthogonal longitudinal field lines.

Assuming that the preferred orientations of OSGCs follow spherical logic, three families of geometrical models could predict the bias toward cardinal orientations in the retina^37,38,41,46,47^. The ‘longitudinal geometry’ model holds that horizontally- and vertically-preferring OSGCs follow two *longitudinal* fields, rotated by 90° from one another (Figure 1C,D); while the ‘latitudinal geometry’ model proposes that OSGCs follow two *latitudinal* fields rotated by 90° from one another (Figure 1E,F). Each of these model families is restricted to a single geometry, either longitudinal (translatory-like) or latitudinal (rotatory-like). A third competing model family blends the two geometries. This ‘hybrid geometry’ model postulates that horizontally-preferring OSGCs follow longitudinal geometry, while vertically-preferring OSGCs follow latitudinal geometry, or vice versa, such that the longitudinal and latitudinal fields are rotated, at any given point, by 90° from one another (Figure 1G,H). All three models adhere to the cardinal orientations in the central retina, and diverge from them in the peripheral retina. However, while all three predict the same cruciform pattern, with a ∼90° angle between preferred orientations in the central retina, only the ‘hybrid-geometry’ model predicts a cruciform pattern, universally, throughout the retina. Interestingly, OSGCs have been recently suggested to topographically match their preferred orientations with concentric patterns^48^, supporting the adherence of OSGCs’ preferences to spherical logic.

In addition to retinal topographical variation in the preferred orientations of OSGCs, the prevalence of OSGCs and/or their OS tuning could also vary topographically across the retina. Retinal topographic variation is common^49^, with variation in cone opsins expression being a fundamental example. The mouse retina includes two cone opsins: a medium (M) wavelength-sensitive opsin peaking at 510 nm (green), and a short (S) wavelength-sensitive opsin peaking at 360 nm in the UV range^50^. However, due to co-expression of the S-opsin in ventral-retina M-cones^51–53^, the ventral retina is especially sensitive to UV light. This retinal topographic variation along the dorsoventral axis may point to a functional specialization, in which the UV-sensitive ventral retina monitors the UV-rich upper visual field to detect predators^54^, while the green-sensitive dorsal retina supports foraging and prey capture^55,56^. Unlike the variation in the expression of cone opsins, topographic variation of OSGCs has been poorly studied.

Here, we tested the correspondence of the three model families with the preferred orientations of large populations of OSGCs across the retina. We identified four OSGC types with different response kinetics. In the central retina, most cells, of each type, preferred horizontal orientation while the rest preferred vertical orientation. However, elsewhere in the retina, the preferred orientations deviated from orthogonality and the cardinal orientations. The preferred orientations of three types could be best explained by the longitudinal geometry model, while those of the fourth type could be best explained by the latitudinal geometry model. Developing a decoder for edge orientation, revealed that, in contrast to the accepted notion, two precisely perpendicular OS sensors would not allow optimal orientation decoding. Instead, optimal decoding is obtained when the angular separation between two OS sensors (corresponding to two OSGCs or two OSGCs ensembles) is the one we found in the longitudinal/latitudinal fields of each type. On the other hand, transforming the OS preferences of cells from *retinocentric* to *egocentric* coordinates, revealed that retinotopic feeding is better served by OSGCs adhering to latitudinal geometry in the monocular zone, while decoding in the binocular zone is best served by vertical-preferring OSGCs adhering to longitudinal geometry and by horizontal-preferring OSGCs adhering to latitudinal geometry. Lastly, OS tuning was narrower in the dorsal vs. ventral retina, mirroring a dorsoventral gradient in the incidence of edges in the natural visual environment of the mouse, and thus may reflect an evolutionary adaptation.

## Results

We mapped response polarity and orientation preference of>33,900 cells at known locations in the ganglion cell layer of flattened mouse retinas *in vitro* (26 retinas, Figure 1I). Cell responses were determined from calcium signals evoked by moving bars and sinusoidal gratings, imaged by two-photon microscopy of virally expressed GCaMP6f (*Methods*). The same dataset was previously used in studying the geometric architecture of DSGCs^44^. Here, response polarity was estimated based on the cell’s response to a bright bar on a dark background drifting perpendicularly to the bar’s long axis in 8 directions (45° intervals), whereas OS was estimated based on the response to sinusoidal gratings drifting in the equivalent 8 directions. While orientation-selective cells are optimally activated by stationary bars projected on their receptive field center ^46^, moving stimuli have been demonstrated to allow detection and identification of these cells^41,43^. Cells were firstly considered orientation-selective if exhibiting an orientation selectivity index (OSI)> 0.2 ^38,43^ and a direction selectivity index (DSI)<0.17 (excludes cells that may be primarily direction-selective)^43,57^, provided that the fit of two Gaussian distributions against the cell’s response amplitude was statistically significant across the 8 tested directions of drifting gratings (*Methods*).

Over 2100 orientation-selective cells were identified, representing 6.2% of all imaged cells (Figure 1J), as presented on a standardized flattened retina and when viewed in a reconstructed three-dimensional form (Figure 1K). Principal component analysis with gaussian mixture model clustering, revealed distinct types of orientation-selective cells, with the optimal number of types being 4. Both the leading and trailing edge of the bar evoked calcium increases in orientation-selective cells belonging to these types, while the cells’ ON and OFF response components varied. The response of one type was dominated by a transient OFF component (‘ON-OFF transient’ or T_ON-OFF_, *n=*620; 1.9% ± 0.6% of imaged cells per retina, mean ± s.d.); the response of a second type was dominated by a sustained OFF component (‘ON-OFF sustained’ or S_ON-OFF_, *n=*273; 0.9% ± 0.3%); the response of a third type was dominated by a transient ON component (‘ON transient’ or T_ON_, *n=*457; 1.4% ± 0.4%); and the calcium signal of the fourth type was dominated by a sustained ON component that remained high until the bar exited the receptive field (‘ON sustained’ or S_ON_; *n=*767; 2.3% ± 0.5%; see *Methods* for bar entry and exit time to and from the receptive field) (Figure 2A). Figure S1A shows responses of individual cells of each type to bar- and grating-motion, with the fit of the grating response amplitude to Gaussian distributions.

**Figure 2.**
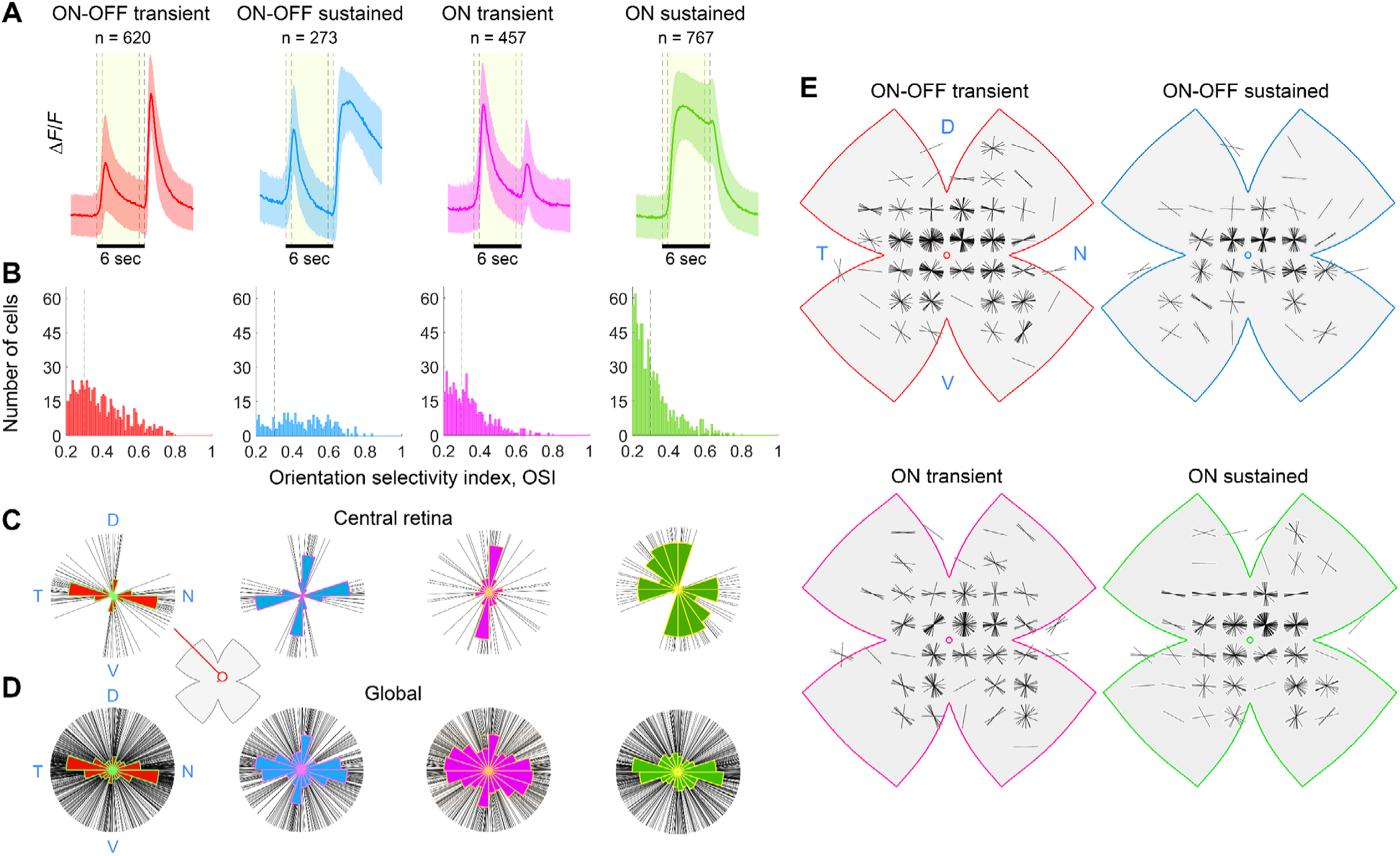
Kinetics, orientation preferences and topography vary across OSGC types. **A.** Calcium responses [Δ*F*/*F*, mean (line)±s.d. (shaded area)] evoked by a light bar moving in the cell’s preferred orientation, for each of the four identified OSGC types: ON-OFF transient (red), ON-OFF sustained (blue), ON transient (magenta), ON sustained (green). Shaded yellow area and dashed black lines mark the approximate time the bar is within the cell’s receptive field. **B.** Distribution of OS index (OSI) for each OSGC type. OSGCs with OSI>0.3 (black dotted line) were included in further analyses. **C, D.** OS preference among imaged OSGCs, one line per cell, pooled from a circled region (marked on a flattened retina inset) in the central retina (**C**), or from the whole retina, for each type. Polar histograms are overlaid. **E.** Topographic dependency of OSGC local OS preferences, displayed as polar plots on a standardized flattened retinal map, for each type.

Evidence for a uniform spatial distribution of OSGCs belonging to each of the four types could support their classification^58,59^. However, such analysis typically requires sampling large, uninterrupted retinal areas, and the imaged fields of view (FOVs) in each retina were rarely adjacent, yielding inconclusive results (Figure S2). Nonetheless, the responses of the four OSGC types to bar motion were consistent across three criteria: (1) OSI > 0.3, (2) OSI > 0.4, and (3) OSI > 0.3 with posterior probability > 0.9 to belong to each of the types (Figure S3), supporting the classification of these functional types.

Six of the 26 imaged retinas were counterstained against the pan-ganglion marker RNA-binding protein with multiple splicing (RBPMS)^60^. Of the orientation-selective cells in those retinas, 88% were RBPMS-positive, and thus were OSGCs (*n=*438). The remaining 12% were distributed across the four functional types, corresponding to a minor fraction of the total cells belonging to each type (Figure S1M-Q). Thus, RBPMS-negative cells are likely poorly RBPMS-labeled OSGCs, but may also correspond to orientation-selective amacrine cells^44^. Cells belonging to each type exhibited a continuum of OSI (ranging from our initial cutoff 0.2, to 0.8). The T_ON-OFF_, T_ON_, and S_ON-OFF_ types, exhibited a smooth right-skewed distribution, while the S_ON-OFF_ type showed a uniform distribution (Figure 2B). To ensure that central-retina OSGCs exhibit preferred orientations roughly aligned with the cardinal orientations, as reported for the mouse and other mammals^37–41,43,46,47,61–64^, and as predicted by the three families of geometrical models (Figure 1C-H), we restricted subsequent analysis to cells with OSI>0.3 (Figure S4). This did not have noticeable effects on the OSGCs’ mean response to bar motion (compare Figure 2A and Figure S3A), and minimized non-OSGCs inclusion in analyses^40,46^.

### Orientation preferences of OSGCs do not globally adhere to cardinal orientations

In all four types, the orientation preferences of OSGCs, pooled across retinas and displayed in polar format, were primarily horizontal, but were markedly disordered (Figure 2D). One or two concentrations of preferred orientations marking the cardinal orientations were apparent, but blurred, except when inspecting only central-retina OSGCs (Figure 2C). This trend was especially salient in the two ON-OSGC types. The disorder formed mainly from systematic topographic variation in OS preferences, as evident through mapping each cell’s location to spherical coordinates, permitting comparisons across retinas using standardized displays as flattened spherical surfaces^44^ (Figure 2E). Away from the central retina, the concentrations of preferred orientations tilted relative to the cardinal orientations and to one another (Figure 2E). Such topographic dependence is not an artefact of retinal flattening, as it is even more apparent when viewed in the reconstructed three-dimensional form (Figure S5A). Thus, plots of OS preference are not ubiquitously cruciform, and do not globally adhere to cardinal orientations.

### Alignment between the OS preferences of OSGCs with three families of spherical models

We tested how well the longitudinal, latitudinal, and hybrid families of geometric models describe the orientation preference of OSGCs and the bias toward cardinal orientations in the central retina. To quantify the degree of alignment between OS preferences per functional type and the *longitudinal* models, we devised a concordance index – the percentage of cells preferring orientations within ±10° of local orientation of the tested longitudinal model (*Methods*). Repeating such template-matching for multiple members of the longitudinal family, we generated a tuning map, displaying concordance as a function of the axis of longitude. Each member (axis) is defined by two coordinates: polar direction (0-350°; 10° resolution) and eccentricity (0-170°, 10° resolution), together defining 684 unique axes. Tuning map hotspots represent the axes forming a longitudinal field best aligned with the observed OS preferences (better concordance), and thus maximal net output of this neuronal ensemble (Figure S5B-D).

For the T_ON-OFF_ type, when the degree of alignment between OS preferences and the *longitudinal* model family was calculated (Figure S5D,E), the full-range tuning map exhibited two identical pairs of hot spots, one within the visual hemifield (lower half of the map) and another outside of it (map upper half). Here and elsewhere, to visualize only one pair of unique hotspots, orientation tuning maps are presented only for the visual hemifield (eccentricity 0-90°) and only for the direction range 120-300° (Figure 3A). For T_ON-OFF_, these two hotspots were separated by about 100° (or the reciprocal 80°) in polar direction (abscissa), suggesting preference for two roughly orthogonal longitudinal axes (Figure 3B). The deviation from orthogonality, which might have intriguing significance, is addressed below. On a flattened retina (Figure 3G), one hotspot is located in the temporal retina; we denote these ‘H-cells’, after their preference for orientations along the nasal-temporal axis in the central retina, and their preference for horizontal edges in the visual field. The other hotspot, denoted ‘V-cells’, is located in the ventral retina; we denote these ‘V-cells’, after their preference for orientations along the ventral-dorsal axis in the central retina, and their preference for vertical edges in the visual field. Therefore, OSGCs of the T_ON-OFF_ type comprise two populations, roughly preferring either horizontal or vertical edges in the visual space.

**Figure 3.**
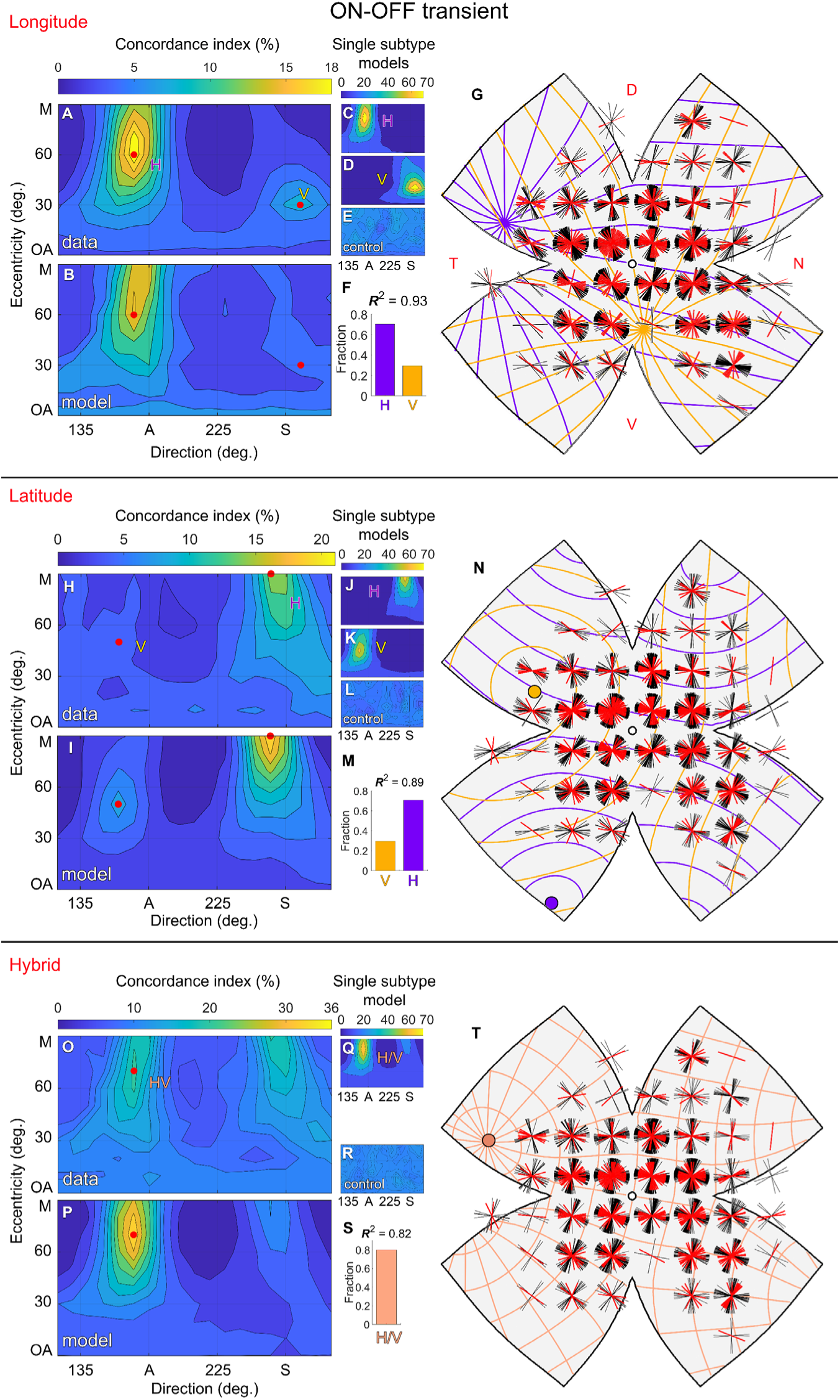
Orientation preferences of T_ON-OFF_ OSGCs align with a longitudinal geometry. **A**. Concordance index as a function of the axis of the longitudinal geometry model, defined by the retinal coordinates: ‘direction’ and ‘eccentricity’, for T_ON-OFF_ OSGCs (n = 425). In this tuning map, eccentricity spans 0° (OA, optic axis) to 90° (M, margin of the visual hemifield, corresponding to the retinal margin). Direction spans 120° to 300°, with direction 180° corresponding to the anterior (A) visual field or temporal retina, and direction 270° corresponding to the superior (S) visual field or ventral retina. Hotspots (red dots) in the anterior and superior visual field represent the locations of axes of the longitudinal model that are best aligned with the OS preferences of T_ON-OFF_ OSGCs. **B-F**. The longitudinal tuning map is recapitulated by a model (**B**) comprising two subtype ensembles: H-cells and V-cells (**C, D**), weighted as in (**F**). **E.** Control longitudinal tuning map created by randomizing OS preferences. **G**. Flattened standardized retina overlaid with a grid of local OS preference polar plots for modelled (black) and real (red) cells. The two best fitting longitudinal axes are represented by purple and yellow field lines. **H-N**. Same as (**A-G**), but for the latitudinal model, with the axes of V- and H-cells reversed (as latitudinal fields are orthogonal to longitudinal fields). **O-T**. Same as (**A-G**), for the hybrid model. One hotspot appears, corresponding to the shared longitudinal-latitudinal axis best aligned with observed OS preferences, denoted ‘H/V-cells’.

To quantify the predictive power of this longitudinal model of OS preferences of T_ON-OFF_ cells, we modelled two T_ON-OFF_ channels adhering to the inferred longitudinal geometry. Each channel comprised modelled cells with locations matched to imaged OSGCs but with preferences aligned exactly with one of the two cardinal longitudinal fields. Angular jitter (about 10°) was added to mimic biological and experimental variability. Each tuning map for single modelled channels exhibited one hotspot (Figure 3C,D). Randomizing modelled preferences yielded a map of uniformly mediocre concordance (Figure 3E). We differentially weighted each channel to reproduce its apparent abundance in our sample and summed them. The tuning map for the best-fitting model (Figure 3B) was strikingly similar to that of the recorded data (Figure 3A; *R*^2^=0.93). The fit was optimal when the relative abundance of H-cells was more than twice that of V-cells (Figure 3F). Local polar plots of OS preference for modelled cells faithfully reproduced those for real cells (Figure 3G).

Next, we compared T_ON-OFF_ OS preferences with the *latitudinal* model family. The latitudinal tuning map also exhibited two hotspots corresponding to H- and V-cells, separated by 100° in polar direction, suggesting again preference for two roughly orthogonal latitudinal axes (Figure 3H). We modelled two T_ON-OFF_ channels adhering to the inferred latitudinal geometry (Figure 3J,K). Fitting the tuning maps for single modelled channels to the data also indicated that the relative abundance of H-cells was over twice that of V-cells (Figure 3M). The fit of the best-fitting latitudinal model, was slightly worse than for the longitudinal model (*R*^2^=0.89; Figure 3H,I,N). Finally, the alignment of the OS preferences of T_ON-OFF_ with the *hybrid* model family was assessed. We generated tuning maps displaying concordance as a function of the shared axis of longitude field and the 90°-rotated latitude field (Figure 3O). The hybrid tuning map exhibited one hotspot, corresponding to the shared longitudinal-latitudinal axis best-aligned with the observed OS preferences. In effect, this axis combines the contribution of H- and V-cells, that prefer either horizontal or vertical edges (hereafter ‘H/V-cells’). Fitting the hybrid geometry T_ON-OFF_ channel to data (Figure 3Q), yielded the combined abundance of H- and V-cells (Figure 3S), with the best-fitting hybrid model (Figure 3P) having worse fit compared to the best-fitting longitudinal and latitudinal models (*R*^2^=0.82; Figure 3O,P,T).

The *R*^2^ confidence intervals (CI) for the longitudinal, latitudinal, and hybrid models (derived using bootstrapping to estimate two-tailed 95% CI), did not overlap, and the *R*^2^ value obtained for the longitudinal model was significantly larger (p>0.05) than those obtained for the latitudinal and hybrid models (Figure 4A). Therefore, the longitudinal model has a statistically-significant advantage in explaining the preferences of T_ON-OFF_ OSGCs.

**Figure 4.**
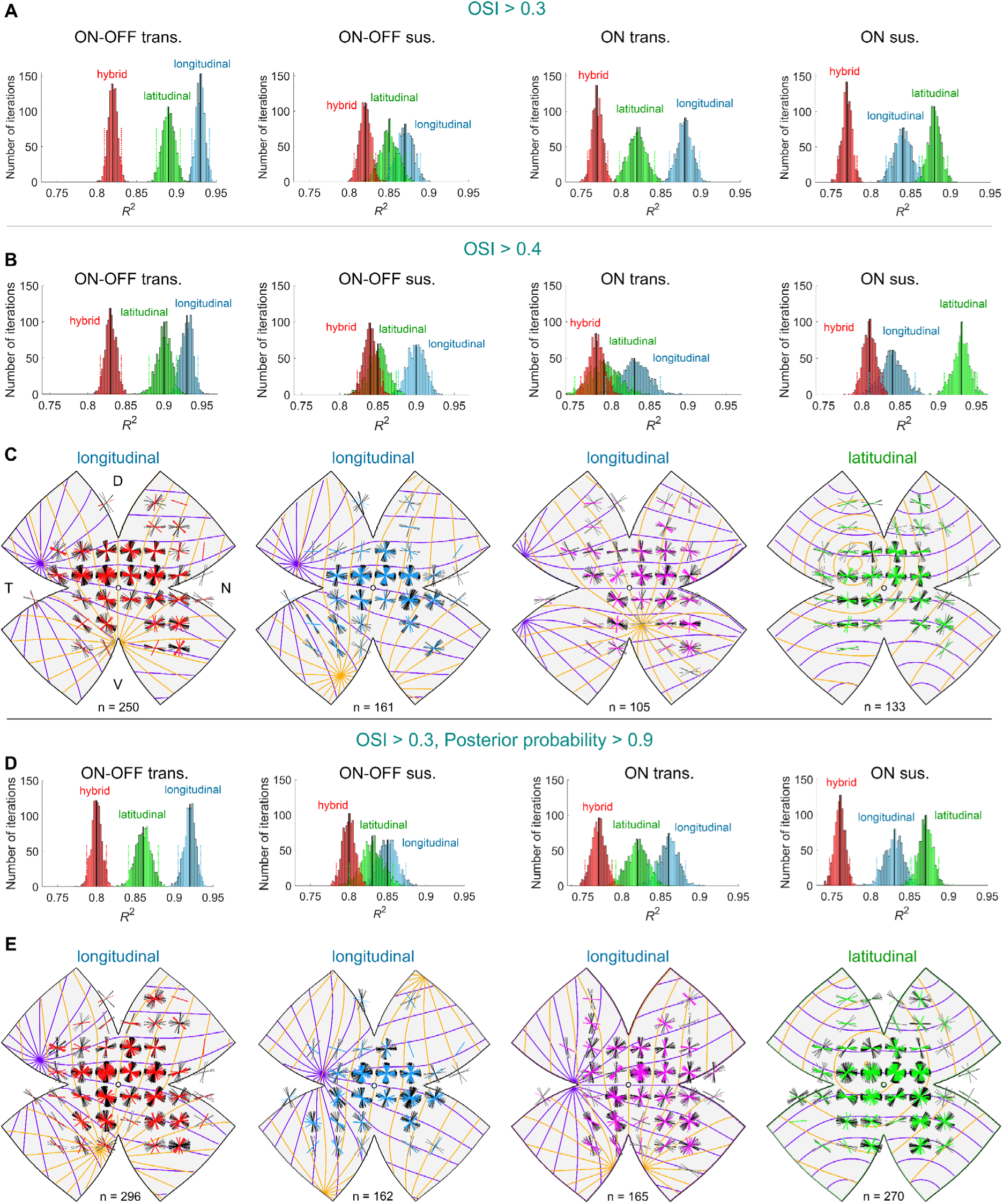
Orientation preferences of S_ON_ align with a latitudinal geometry while those of remaining types align with a longitudinal geometry. **A.** Significance testing of the alignment with the longitudinal (blue), latitudinal (green) and hybrid (red) geometric models, of the OS preference of cells with OSI>0.3 for the four OSGC types, using bootstrapping on *R*^2^ values. Black lines, median; dotted colored lines, *R*^2^ confidence interval (CI). The longitudinal model was superior in describing the OS preference of T_ON-OFF_, S_ON-OFF_, and T_ON,_ while the latitudinal model was superior in describing the OS preference of S_ON_ (T_ON-OFF_ – CI_long_: 0.908 - 0.930, CI_lat_: 0.874 - 0.904, CI_hybrid_: 0.809 – 0.830; S_ON-OFF_ – CI_longitudinal_: 0.850 - 0.889, CI_latitudinal_: 0.829 - 0.870, CI_hybrid_: 0.806 – 0.833; T_ON_ – CI_longitudinal_: 0.861 - 0.899, CI_latitudinal_: 0.797 - 0.842, CI_hybrid_: 0.758 – 0.783; S_ON_ – CI_longitudinal_: 0.819 - 0.861, CI_latitudinal_: 0.864 - 0.896, CI_hybrid_: 0.758 – 0.783). **B.** Alignment testing for cells with OSI>0.4. **C.** Flattened standardized retinas overlaid with a grid of local OS preference polar plots for real cells with OSI>0.4 (colored) and cells modelled by the best fitting longitudinal, latitudinal or hybrid geometry (black). **D,E.** The same as (**B,C**) for cells with a posterior probability>0.9 to belong to each of the types (as opposed to the default posterior probability>0.5).

We repeated the same procedures for the remaining three OSGC types (Figure S6 and S7). As in the case of the T_ON-OFF_, the hybrid model poorly explained the variation in OS preferences in the remaining three OSGC types. For the S_ON-OFF_ and T_ON_ types, the longitudinal model was superior in describing OS preference, albeit not in a statistically significant manner (Figure 4A). However, surprisingly, the OS preference of the S_ON_ type could be best predicted by the latitudinal model (Figure 4A). The same favorable explanatory power of longitudinal geometry for the T_ON-OFF_, S_ON-OFF_, and T_ON_ types and of latitudinal geometry for the S_ON_ type was obtained when repeating the analysis only for cells with OSI>0.4 (Figure 4B,C), or only for cells with a posterior probability>0.9 to belong to each of the types (as opposed to the default posterior probability of>0.5) (Figure 4D,E).

In all types, H-cells accounted for 56-72% of all OSGCs, when modelled with either the longitudinal or latitudinal models (Figures 3F, S5C, S6C, and S6T). The three types best aligned with a longitudinal geometry varied slightly in their best axes, and the ‘direction’ coordinates of those best axes averaged at 173° and 273° for OSI>0.3 and OSI>0.4 criteria (Figure 5A). These best axes roughly match the translatory best axes identified previously in DSGCs (185° and 275°)^44^. However, critically, the best axes of OSGCs were displaced along the direction axis by 80° (or the reciprocal 100°), but those of DSGCs were displaced by 90°. Likewise, the ‘direction’ coordinates of the best axes of the S_ON_ type, which was best aligned with a latitudinal geometry, averaged at 135° and 270° (Figure 5A), representing a displacement by 135° (or the reciprocal 45°) along the direction coordinate. That is, the displacement of the best axes along the direction coordinate of each of the four OSGC types deviates from orthogonality.

**Figure 5.**
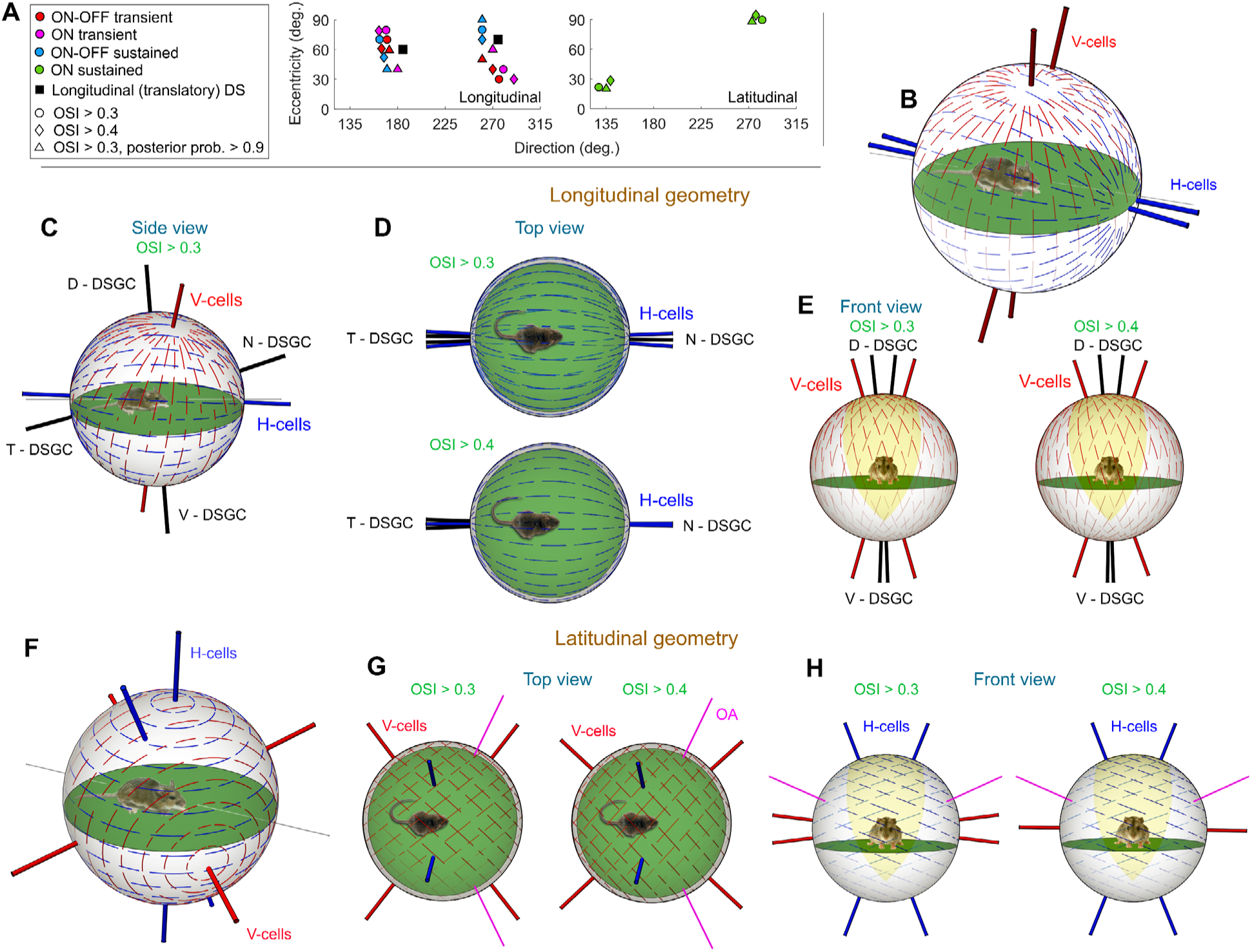
Retinal OS ensemble-coding of horizontal and vertical edges is consistent with longitudinal and latitudinal geometry. **A.** Retinal coordinates of best-aligned axes (hotspots) for T_ON-OFF_, T_ON_, and S_ON-OFF_ (adhering to a longitudinal geometry), and for S_ON_ (adhering to a latitudinal geometry). Custering of the best axes is greater along the direction-coordinate than along the eccentricity-coordinate. Black squares, axes of two translatory (longitudinal-like) optic flow patterns best aligned with the direction preferences of DSGCs. **B-E.** The longitudinal geometry. **B.** The longitudinal axes of H- and V-cells of both eyes, and the resulting fields in visual space (presented only for the right eye). The axes of _long_H-cells (blue) and _long_V-cells (red) approximated the anteroposterior (body) axis and the gravitational axis, respectively. **C.** Side view, best axes and fields of the right eye’s _long_H-cells (blue) and _long_V-cells (red); best axes of DSGCs are depicted in black. **D,E.** Best axes and fields of _long_H-cells (**D**, blue, top view) and _long_V-cells (**E**, red, front view) in both eyes, for cells with OSI>0.3 or OSI>0.4; best axes of DSGCs are depicted in black. **F.** latitudinal axes of _lat_H-cells (blue) and _long_V-cells (red) of both eyes, and the resulting fields in visual space (presented only for the right eye). **G,H.** Best axes and fields of _lat_V-cells (red) in both eyes, top view (**G**), and of _lat_H-cells (blue) in both eyes, front view (**H**). In both panels, axes and fields are presented for cells with OSI>0.3 (left) and OSI>0.4 (right).

We repeated the same analysis while estimating the OSI and preferred orientations of OSGCs based on their responses to bar instead of gratings motion. OSI based on bar motion, accounting for the ON and OFF response components, was lower than that obtained from gratings motion, indicating that drifting gratings better engage OSGCs (permutation paired t-test; p < 0.001 for all types; T_ON-OFF_, *n=*418, OSI_grating_ = 0.38 ± 0.13, OSI_bar_ = 0.14 ± 0.10; S_ON-OFF_, *n=*220, OSI_grating_ = 0.45 ± 0.15, OSI_bar_ = 0.23 ± 0.13; T_ON_, *n=*259, OSI_grating_ = 0.34 ± 0.11, OSI_bar_ = 0.10 ± 0.07; S_ON_, *n=*322, OSI_grating_ = 0.32 ± 0.10, OSI_bar_ = 0.07 ± 0.06). OS preferences based on either the ON or OFF response components to bar motion varied slightly from those based on grating motion, diverging up to 60° for H-cells in S_ON_ but usually within 10°-20° (Figure S8). The relative weighting of H-cells and V-cells in the best-fitting longitudinal or latitudinal model was nearly identical for ON and OFF response components to bar motion, and for grating motion (Figure S8). Therefore, while drifting bars yield lower OSI values than gratings, OSGC OS preferences and spherical alignment remain largely similar.

### OSGCs encoding of edge orientation follows two distinct spherical geometries

We converted the best longitudinal/latitudinal axes from a retinal space (retinocentric) to an extra-personal space (egocentric), relative to the visual environment, while accounting for the resting eye and head positions (*Methods*). To facilitate a comparison between a longitudinal and latitudinal geometry, a pair of OS longitudinal best axes (_long_H-cells and _long_V-cells) was determined based on the mean of the axes of the three OSGC types adhering to longitudinal geometry (T_ON-OFF_, S_ON-OFF_, and T_ON_ types), and a pair of OS latitudinal best axes (_lat_H-cells and _lat_V-cells) were taken as the axes of the S_ON_ type that adheres to latitudinal geometry.

The *longitudinal* axis of OS H-cells (_long_H-cells) approximated the anteroposterior (body) axis, whereas that of V-cells (_long_V-cells) roughly corresponded to the gravitational axis (Figure 5B). Thus, while adhering to longitudinal geometry, OS _long_H- and _long_V-cells, *as ensembles*, roughly respond best to global horizontal and vertical edge orientations. This is reminiscent of the DSGC N-, T-, V-, and D-cells, whose preferred directions are best aligned with the optic flow patterns casted on the retina when the animal moves forward and backwards along the body axis (N- and T-DSGCs), and upward and downward along the gravitational axis (V- and D-DSGCs)^44^; however, as a side view of the coordinate system reveals, the OS longitudinal axes are rotated by ∼20° relative to DS translatory axes (Figure 5C). Moreover, the axes of _long_H-cells of the two eyes diverge by 6.8° when cells with OSI>0.3 are considered, and when only cells with OSI>0.4 are considered become virtually identical (angle difference=1.4°), as a top view of the coordinate system reveals (Figure 5D). Therefore, _long_H-cells, *as an ensemble*, prefer the same edge orientation regardless of their location in either retina. In contrast, the axes of _long_OS V-cells of the two eyes diverge by 32° with OSI>0.3 and by 37° with OSI>0.4, and thus _long_V-cells, *as an ensemble*, prefer a different edge orientation in each eye, as a front view of the coordinate system reveals; this divergence was larger than that in the axes of D/V-DSGCs (Figure 5E). The *latitudinal* H-cells and V-cells (_lat_H- and _lat_V-cells), *as ensembles*, also responded best to global horizontal and vertical edge orientations (Figure 5F). However, the axes of _lat_H-cells were tilted (22°) from the gravitational axis, while those of _lat_V-cells were approximately aligned with the horizontal plane and deviated from the optic axes by 23° (Figure 5G-H). Therefore, _lat_H- and _lat_V-cells, *as ensembles*, prefer a different edge orientation in each eye.

### Deviation from orthogonality between OS sensors enables efficient decoding of edge orientation

OS in the visual cortex is thought to be important for perception of edges’ orientation^3,4,7–15,17^, whereas retinal OS may contribute to additional functions. These include the detection of the horizon, maintenance of head and body orientation, localization of objects in the visual field, and regulation of defensive responses^65,66^. While such functions may not require precise edge orientation decoding, they would likely benefit from it or remain unaffected. Thus, we sought to explore the ability of retinal OS to enable precise edge orientation decoding by post-synaptic neurons.

The central brain targets of OSGCs likely include the SC and dLGN – both exhibiting OS^27–30,34,67–69^. OSGCs may feed neurons in those and other brain targets in two general configurations: (1) as an ensemble, with a postsynaptic neuron integrating signals from multiple OSGCs across the retina, or (2) retinotopically, with minimal integration from OSGCs in different retinal locations. A postsynaptic neuron (either a retinorecipient or higher-order neuron) fed by two OS sensors, *e.g.*, H- and V-cells, either retinotopically or as an ensemble, might constitute an edge-orientation decoder. However, biological noise in the orientation preference of OSGCs would presumably degrade the efficiency of such a decoder. To determine the angular separation between two OS sensors required for efficient edge orientation decoding, we constructed a geometrical decoder that accounted for the biological noise in OSGCs’ preferred orientation (4.457°, *Methods*). See Figure S9 and *Supplementary equations* for a detailed description of the development of the decoder.

The preferred orientation of each sensor was modeled using the von Mises probability density with its standard deviation set to the biological noise in the preferred orientation of OSGCs. The response evoked by a real-world edge (*E*) was modeled using the cosine of the angle formed between each sensor’s preferred orientation and the orientation of the real-world edge (Figure S9A). The modeled response of each sensor was found to be bimodal, being concentrated about the real edge orientation (*E*) and an ‘illusionary edge’ (*E*^∗^), with the two forming the same angle with the sensor’s preferred orientation. The illusionary edge results from the fact that the reflected image of any edge about the preferred orientation of any sensor yields the same projection onto the sensor. Relying on information from two sensors would thus increase the probability of decoding the real edge rather than an illusionary one. Figure S9B-D illustrates relative positions of the real and illusionary edges resulting from different angles between the two sensors.

To decode the orientation of a real-world edge, *i.e.*, to use the responses of two sensors to determine the edge’s orientation, we inverted the encoding procedure. This yielded, for a given real-world edge’s orientation, a probability density representing the distribution of decoded edge orientations (*υ*) from the responses of two sensors (Figure S9E,F). Repeating this procedure for a range of real-world edge orientations resulted in a 4-dimentional matrix representing the probability density as a function of the angle between the mean preferred orientations of the sensors (*ϕ*), the angles between the real-world edge and the preferred orientation of the sensors (*θ* and *ϕ* − *θ*), and *υ* (Figure S9G, Video S1). The efficiency of edge orientation decoding (*δ*) was taken as the difference in probability of decoding the orientation of the real edge vs. the illusionary edge, across all the possible real-world edge orientations. Efficient orientation decoding (*δ*>0.95) occurred with the angle between the sensors (*ϕ*) being larger than 8.5° and smaller than 81.5°, regardless of the orientation of the real-world edge (Figure S9H). Therefore, in contrast to the accepted notion, two orthogonal OS sensors would *not* constitute the optimal decoder. Such a decoder would confound edges with orientations symmetric about the sensors. In fact, only a decoder fed by two sensors that deviate from orthogonality would allow the discrimination between such orientations. Additionally, biological noise in the orientation preference of OSGCs would degrade decoding efficiency – the smaller the angle between sensors, the lower the decoding efficiency would be. In summary, either a too small (<8.5°) angle between the sensors, or too small a deviation of the two sensors from orthogonality (angle between sensors>81.5°), would degrade decoding efficiency. These predictions of decoding efficiency are independent of the actual postsynaptic input convergence in the brain region in question, i.e., whether postsynaptic neurons recieve inputs from ensembles of OSGCs or retinotopically from one or a few OSGCs.

However, remarkably, for an ensembled operation (configuration #1, above), the slight deviation from orthogonality (80°) that we discovered between the preference of H- and V-cells (Figure 5A), also evident in the preferences of central retina OSGCs (Figure 2C,D), would enable efficient orientation decoding. While the currently known brain targets of OSGCs (SC and dLGN) exhibit retinotopic organization^27–30,34,67–69^, other brain regions, yet unidentified, may receive input from OSGCs ensembles. In that case, the OS preference of H- and V-cells would allow close to optimal decoding of edge orientation.

### The decoding efficiency of longitudinal and latitudinal OSGCs varies across the monocular and binocular zones

The case of OSGCs feeding their brain targets retinotopically (configuration #2) is more complex, as decoding efficiency would depend on the angular disparity between neighboring H- and V-cells, which varies topographically across the retina, and between visual field-topographically-matching H-cells or V-cells from both eyes. To identify visual field regions in which the angular disparity between two OS sensors would allow efficient edge-orientation decoding, we calculated the angular separation between key pairs of OS sensors (_long_H-cells vs. _long_V-cells of the right eye; _long_H-cells right vs. left eye; _long_V-cells right vs. left eye), and estimated decoding efficiency across the visual field.

#### Monocular zone decoding efficiency maximization by long. vs. lat geometry

Efficiency when relying on _long_H/V-cells or on _lat_H/V-cells of a single eye, was generally high, but exhibited two bands of low efficiency, traversing the monocular and binocular zones (Figure 6A-D). While efficiency varies topographically, to uncover general phenomena we compared efficiency across the monocular and binocular zones by pooling all visual field locations in each zone. In the monocular zone, efficiency was higher when relying on _lat_H/V-cells vs. _long_H/V-cells (Figure 6M).

**Figure 6.**
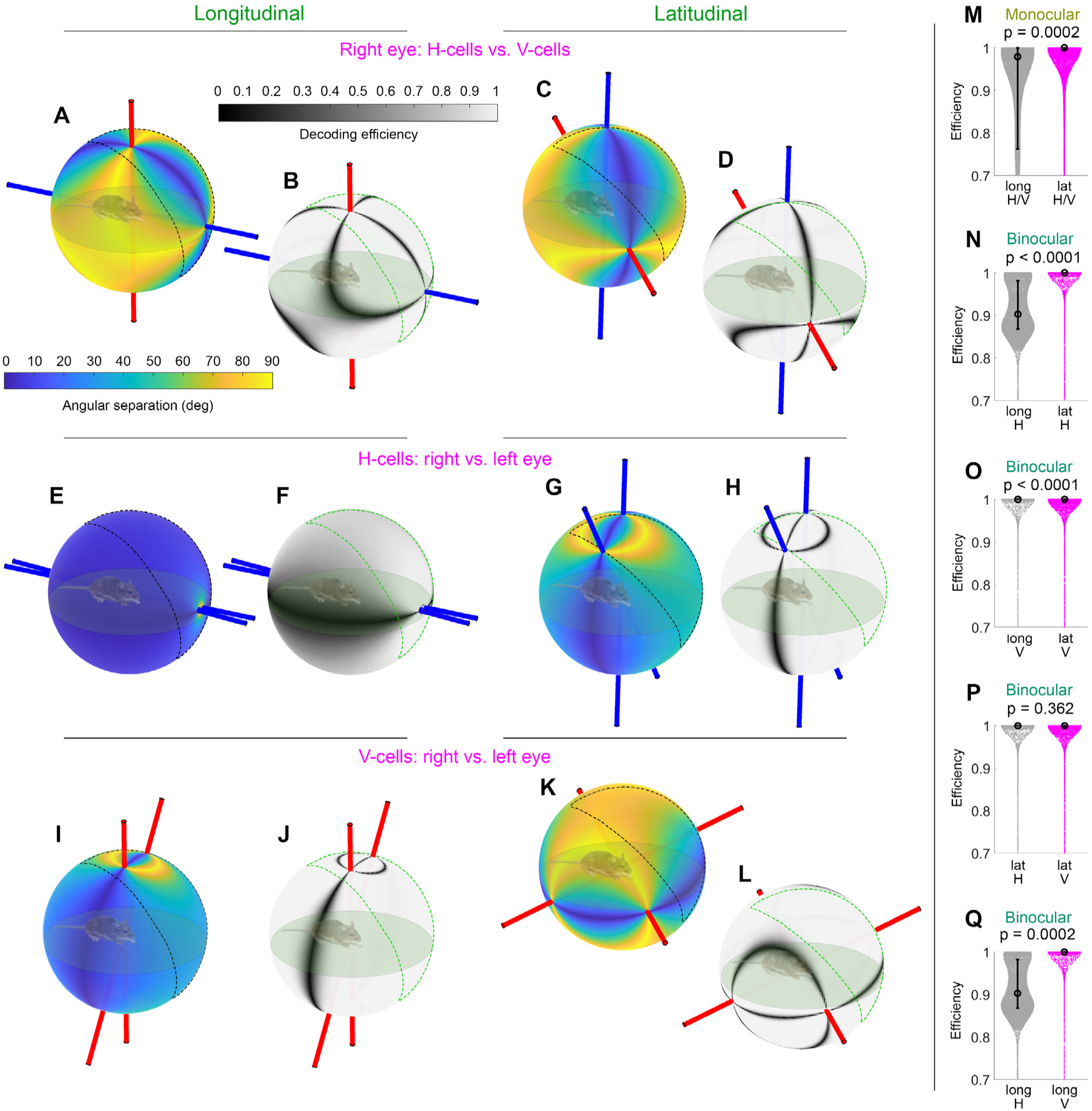
Angular separation and decoding efficiency in the monocular and binocular zones. **A-D.** Angular separation (**A,C**) and decoding efficiency (**B,D**) of _long_H/V-cells and _lat_H/V-cells, right eye. **E-H.** Angular separation (**E,G**) and decoding efficiency (**F,H**) of _long_H-cells in the right vs. left eye and of _lat_H-cells in the right vs. left eye. **I-L.** Angular separation (**I,K**) and decoding efficiency (**J,L**) of _long_V-cells in the right vs. left eye and of _lat_V-cells right vs. left eye. **M,N,** In the monocular zone, decoding efficiency was higher for _lat_H/V-cells vs. _long_H/V-cells (**M**) (permutation t-test, p=0.0002, *n=*46400), while in the binocular zone it was higher for _lat_H-cells of both eyes vs. _long_H-cells of both eyes **(N)** (permutation t-test, p<0.0002, *n=*18941). **O-Q.** In the binocular zone, decoding efficiency was higher for _long_V-cells of both eyes vs. _lat_V-cells of both eyes **(O)** (permutation t-test, p<0.0002, *n=*18941), while not differing between _lat_H-cells and _lat_V-cells of both eyes **(P)** (permutation t-test, p=0.362, *n=*18941), and being higher for _long_V-cells vs. _long_H-cells **(Q)** (permutation t-test, p<0.0002, *n=*18941).

#### Binocular zone decoding efficiency maximization by long. vs. lat. geometry and H-vs. V-cells

Several pairs of OS sensors would allow decoding edge orientation in the binocular zone. We thus calculated the angular separation and decoding efficiency for two key pairs of OS sensors: _long_H-cells right vs. left eye (Figure 6E-H), and _long_V-cells right vs. left eye (Figure 6I-L). Decoding efficiency was higher when relying in both eyes on _lat_H-cells vs. _long_H-cells (Figure 6N), or in both eyes on _long_V-cells vs. _lat_V-cells (Figure 6O). While efficiency was similar when relying on _lat_H-cells or _lat_V-cells of both eyes (Figure 6P), it was higher when relying on _long_V-cells vs. _long_H-cells of both eyes (Figure 6Q).

Thus, in the monocular zone, _lat_H/V-cells, and less so _long_H/V-cells, allow high decoding efficiency of edge orientation, whereas in the binocular zone, either _long_V-cells or _lat_H-cells of both eyes allow the highest efficiency. Constructing the geometrical decoder while increasing the biological noise in OSGCs’ preferred orientation decreased the decoding efficiency (Figure S9I), but did not alter the relationships between decoding efficiency when relying, for instance, on H/V-cells in the monocular/binocular zones (Figure S9J).

### Variation in orientation selectivity across the retina may mirror variation in the statistics of the mouse natural habitat

In all OSGC types, a gradient of OSI across the retina was observed (Figure 7A), with OSI in the dorsal retina being significantly higher than in the ventral retina, for all types together (permutation t-test, p<0.0001, effect size=0.485) and for each type separately (permutation t-test, p=0.0001, 0.0001, 0.0067, and 0.0013 for T_ON-OFF_, S_ON-OFF_, T_ON_ and S_ON_) (Figure 7B,C). The higher OSI in the dorsal retina may represent an adaptation to a greater incidence of edges in the ventral visual field. We utilized a published dataset of footage from natural habitats of mice^70^, that spanned a solid angle of ∼180°, captured from a few centimeters above the ground using a camera maximally sensitive to two spectral bands covering the absorbance spectra of UV and M (green) cone opsins in the mouse. The camera’s optic axis was positioned such that the horizon bisected the captured footage into a lower and upper visual field. Two image crops from each image’s upper and lower halves (128×128 pixels, equivalent to a solid angle of 53°) were extracted. To avoid confounding effects of overexposure on the detection of edges, we analyzed 1000 randomly-sampled crops with mean low and intermediate intensities (*Methods*).

**Figure 7.**
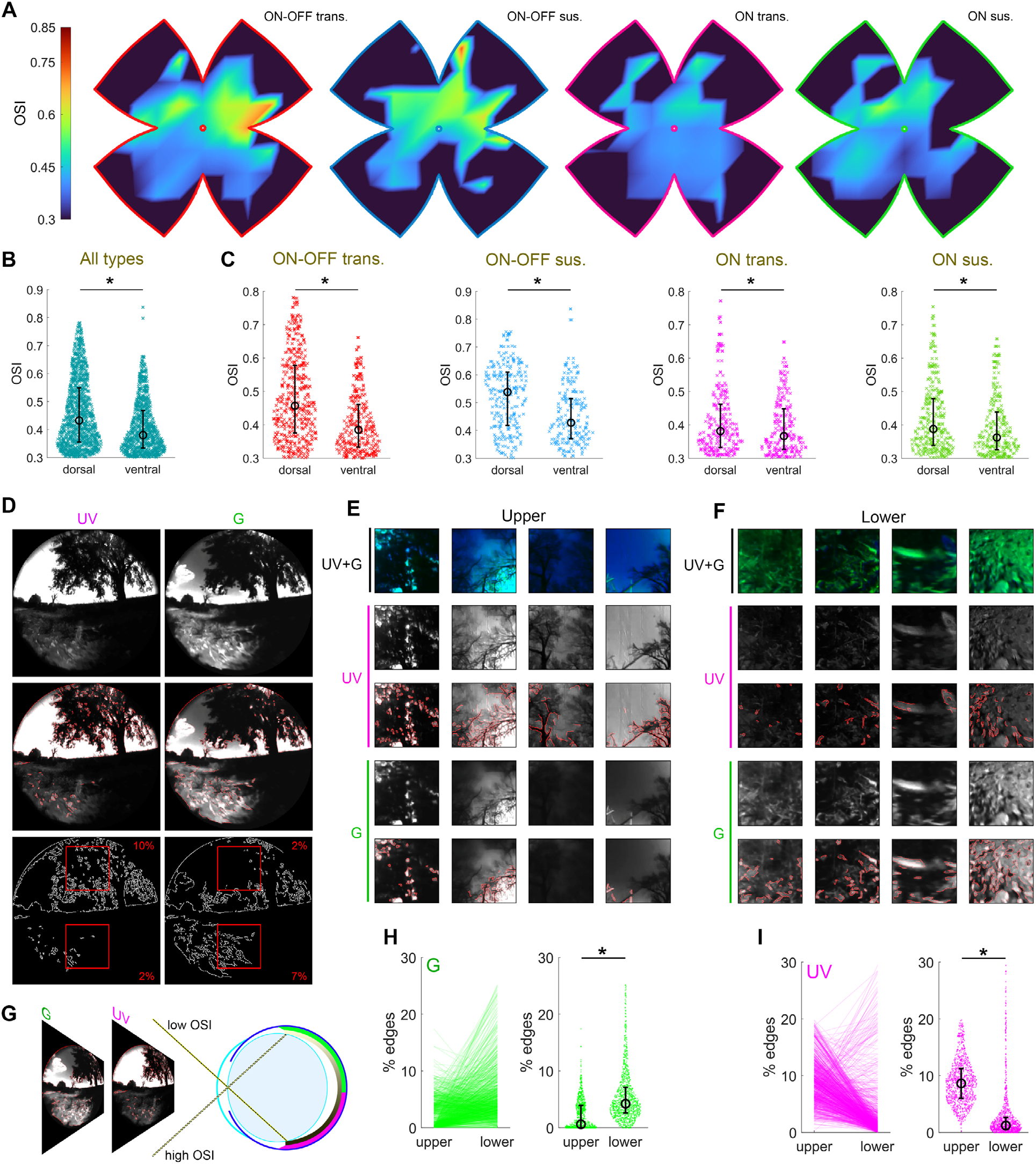
Orientation selectivity changes gradually across the retina. **A.** A heatmap of the mean OSI, ranging from 0.3 (cold) to 0.85 (warm) across the retina, on a computationally-flattened retina, for each OSGC type. **B,C.** OSI in the dorsal vs. ventral retina, for all types (**B**) and for each individual type (**C**). **D.** An example frame, taken from a mouse-habitat movie in the UV and green ‘G’ channels (**top**), with detected edges overlaid (**middle**), and with the detected edges alone (**bottom**). **E,F.** Example image crop in the UV and green ‘G’ channels, presented as a merge of UV+G in the top panels, and individually with and without the detected edges, for the upper (**E**) and lower (**F**) visual space. **G.** Schematic of a retina exhibiting an OSI gradient (grayscale) and a UV sensitivity gradient (UV-green) along the dorso-ventral axis. **H,I.** Incidence of edges in the lower vs. upper halves of the green (**H**) and UV (**I**) channels.

The detection of edges by the retina would depend on the spatial contrast of the retinal image, which is shaped by various factor, including the pupil size, spectral filtering and contrast degradation by the eye optics, as well as retinal density of photoreceptors and other retinal neurons^52,71–77^. Here, for simplicity, the spatial contrast of the retinal image was assumed to be similar to that of the captured image. We developed an algorithm for the detection of edges in the selected crops (Figure 7D). Edges in the UV and green image channels were detected using the Canny Edge Detection^78^ as implemented for Python in the OpenCV library^79^. Figure 7E,F shows example image crops, taken from Video S2, in the two channels, individually and merged, and with the detected edges, for the upper and lower visual space. The incidence of edges was higher in the lower visual field in the green channel, but higher in the upper visual field in the UV channel (permutation paired t-test, p<0.0001 for UV and green, *n=*1000) (Figure 7H,I). This greater incidence of green-channel edges in the lower visual field coincides with the greater incidence of OSGCs and the higher OSI observed in the dorsal retina, while the greater incidence of UV-channel edges in the upper visual field coincides with the higher expression of UV cone opsin reported in the ventral retina^51–53^. Taken together, the retinal topographic variation in OSI and UV cone opsin may represent an adaptation for the detection of edges in the mouse’s natural habitat.

## Discussion

### Orientation preferences of OSGCs follow spherical geometry

The preferred orientations in the central retina of all four identified OSGC types roughly matched the two cardinal orientations, more than elsewhere in the retina. This central-to-peripheral divergence may reconcile discrepancies between reports on variation in OSGCs’ preferred orientations^37–42,46,47^. Orientation preferences of T_ON-OFF_, S_ON-OFF_, and T_ON_ were best aligned with longitudinal geometry, while those of S_ON_ with latitudinal geometry. The bias toward cardinal orientations in the dLGN and V1, observed in response to wide-field stimuli^27–29,80,81^, could be readily explained by retinal drive from OSGCs with orientation preferences following spherical logic – a requirement that all the four OSGC types fulfilled. Independently of their adherence to longitudinal or latitudinal geometry, each of the four types constituted a population of H-cells (preferring roughly horizontal real-world edges) accounting for 56-72% of OSGCs, and a population of V-cells (preferring roughly vertical real-world edges).

The stationary longitudinal and latitudinal models represent a snapshot of the optic flow generated when the animal translates along (longitude), or rotates about (latitude) a given axis. The preferred direction of DSGCs were reported to match two translatory optic flow fields – encoding the animal’s self-motion along the body axis and the gravitational axis^44^. Here, the longitudinal geometric model could explain the preferred orientations of OSGCs of the types T_ON-OFF_, S_ON-OFF_, and T_ON_, but not as good as a translatory optic flow model explained the preferred directions of DSGCs (OSGCs: *R*^2^=0.87-0.93; DSGCs: *R*^2^=0.95-0.97) ^44^. The slightly poorer fits obtained for OSGCs might suggest greater natural variability in the orientation preferences of OSGCs than in the direction preference of DSGCs. Indeed, restricting our analysis to cells with OSI>0.4, typically accentuated the difference in the goodness of fit of OSGCs to the longitudinal vs. latitudinal models.

Our results of the topographical variation of OSGCs’ preferred orientations generally agree with those recently reported in Vita et al^48^. First, the best axes for DSGCs did not align with those of OSGCs, supporting that DS and OS maps are distinct. Second, the majority of OSGCs in the ventro-nasal retina preferred orientations along the ventro-temporal to dorso-nasal axis, whereas most OSGCs in the ventro-temporal retina preferred orientations along the ventro-nasal to dorso-temporal axis. This topographical variation in orientation preferences was observed across all OSGCs collectively and within each of the four identified OSGC subtypes (Figure S10). However, the two studies differ in a key aspect – while Vita et al^48^ suggested that OSGCs’ orientation preferences align with 2-dimentional (2D) concentric ellipses on the flattened retina, the models proposed here are spherical (3D), accounting for the projection of visual space onto the hemispherical retina and for distortions in the location and preferences of cells during the flattening of the retina. Similar models have been shown to succefully predict the preferred directions of DSGCs^44^. In analogy, similar to how 2D geometry can be used for mapping small geographic areas as a local approximation of Earth’s spherical shape, 2D ellipses on a flattened retina approximate its spherical geometry and are suitable only for mapping small retinal patches. Thus, while the 2D geometry model predicts orientation preferences near the ellipse’s center, it fails for cells farther away. Indeed, shifting the ellipses center toward the ventral retina, where OS cells were studied, improved the model’s alignment with the Vita et al^48^ data.

Nath and Schwartz^38^ mapped the location and orientation preferences of ON-OSGCs. We extracted these data, mapped them onto a standard retina, and tested their alignment with the best-fit latitudinal axes identified for the S_ON_ type in the current study. These axes poorly explained the topographical variation in the orientation preferences of ON-OSGCs (*R²* = 0.25; Figure S10C). Testing other latitudinal and longitudinal geometry models also yielded relatively low explanatory power (*R²* = 0.64 and *R²* = 0.75, respectively; Figure S10D,E). The poor fit of Nath and Schwartz’s data with ours or with spherical geometry is unsurprising. Unlike previous studies, we documented each retina’s dimensions, the relieving cuts made during flattening, and the resulting distortions in cell locations and orientations. We then fine-registered the retinas and mapped the cells onto a standard retina, enabling data pooling across all retinas. We argue that precise mapping is essential for studying the global geometry of OS preferences in OSGCs, as of DS preferences in DSGCs^44^.

### OSGCs allow for edge orientation decoding in multiple connectivity schemes

OSGCs might feed their brain targets as ensembles, integrating signals from across the retina, or retinotopically, with minimal integration. Weighted combinations of the two configurations might also exist. Constructing a geometrical decoder, simulating a postsynaptic brain neuron fed by two OS sensors, revealed that for optimal orientation decoding, the deviation between the orientation preference of the two sensors should be between 8.5° and 81.5°.

While a small angular separation between sensors was expected to yield low decoding efficiency, due to biological noise, angular separation approaching orthogonality generated a surprisingly low decoding efficiency. That is, in contrast to the accepted assumption, that OSGCs preferences exhibit orthogonality, two orthogonal OS sensors would *not* constitute the optimal decoder. Instead, and as the decoder modelling predicts, two lines of evidence in our results show that the orientation preference of H- and V-cells deviate from orthogonality: (1) the best axes of H- and V-cells deviate by 80° rather than by 90° along the direction axis, and (2) the orientation preferences of central-retina OSGCs form two lobes deviating from orthogonality. Both trends appear in all four types. Such deviation from orthogonality may represent a specialization of H- and V-cells allowing efficient decoding of edge orientation in an ensembled operation.

Our results further demonstrate that the orientation preferences of OSGCs might also underlie efficient edge orientation decoding in retinotopically-organized operation. In this case, decoding efficiency depends on the angular separation between the subsets of OS sensors available in the monocular or binocular zones. We found that in the monocular zone, _long_H- and _long_V-cells, and even more so _lat_H- and _lat_V-cells, allow high decoding efficiency of edge orientation. Whereas, in the binocular zone, either _long_V-cells or _lat_H-cells of both eyes allow the highest decoding efficiency. This offers specific and testable hypotheses regarding the OSGC types that may underlie a retinotopic edge orientation decoding, in different brain regions, and for different visual field regions. For example, retinal axons’ buttons in the binocular zone of the SC retinotopic map have been shown to prefer orientations biased toward the horizontal^34^. Our results indicate the greater likelihood of _lat_H-cells vs. _long_H-cells to contribute to the formation of the SC binocular retinotopic map.

A brain neuron receiving retinal OS signals, may be a retinorecipient neuron, fed directly by the two OS sensors (H- **and** V-cells), or instead a higher-order neuron fed by two retinorecipient neurons, each innervated by a single OS sensor (either a H- **or** V-cell). Thus, the interface between the signals of the two OS sensors may be manifested at the retinorecipient neuron or downstream. These two possibilities are largely consistent with the two main axon convergence modes reported previously in retinorecipient dLGN neurons: ‘relay mode’ cells integrating inputs from RGCs of mostly one type, and ‘combination mode’ cells combining inputs from many RGC types^82–86^. Interestingly, support for the first possibility, i.e., a retinorecipient neuron fed directly by two OS sensors, comes from studying the fine-scale convergence of retinal input in the dLGN, revealing that dLGN neurons can be innervated by RGC types belonging to the same category (e.g., DSGCs) but with opposite direction preferences^86^. Further support may arrive from a recent report showing that a single DSGC can be innervated by bipolar cells’ axonal buttons with diverse direction preferences^87^. Moreover, instead of decoding based on two sensors (e.g., V- and H-cells), a single OSGC, either a V- or H-cell, might suffice for crude post-synaptic orientation decoding. However, considering the potential confusion between the real and virtual edges, such a configuration would likely enable no more than mere distinction between vertical and horizontal orientations. Importantly, while it is currently unknown whether precise post-synaptic orientation decoding based on retinal OS is manifested or even needed, this study demonstrates that such OSGC-based decoding of edge orientation is certainly possible.

### OSGC diversity

Each OSGC type was found to have a distinct response polarity. Additionally, because we excluded from analysis cells with DSI<0.17, none of the four types likely corresponds to a recently described DSGC receiving both OS and DS inputs^57^. Three of the four OSGC types identified here (S_ON-OFF_, S_ON,_ and T_ON_) fit well with those recently identified using a large dataset obtained though Ca^2+^ in the ventral mouse retina, and being referred to as OFF, ONt, and ONs^48^. The identification of the T_ON-OFF_ type here but not in the previous study may be explained by our observation that these cells exhibit low density and low OSI in the ventral retina, where imaging in the previous study was restricted to. The absence of the T_ON-OFF_ type in the previous study might also be explained by differences in the stimulus used for the calculation of the OSI – drifting sinusoidal gratings here vs. drifting bars in the other study.

Additionally, we propose that cells of the T_ON-OFF_ type detected here but not in the recent Ca^2+^ study may be analogous to the F-mini-ON cells^88,89^. Similarly to our T_ON-OFF_, the F-mini-ON responds both to the leading and trailing edges of a drifting bar^89^. F-mini-ON cells, stimulated by drifting bars and gratings at similar speeds to those we used, have been shown to exhibit low DSI in response to drifting bars, and slightly higher OSI in response to drifting gratings (no OSI was reported for bars, and no DSI was reported for gratings)^89^.

S_ON_ may also correspond to the rabbit horizontally-preferring ON-OSGC^37,46,47^, to the mouse G30 ON-sustained-OSGC^42^, or to ON-OS SmRF, the latter exhibiting light-evoked sustained firing, prefering either vertical or horizontal orientation^41^. T_ON_ could also correspond to G17 ON transient-OSGC (both show high variability in OS preferences)^42^, or to ON-OS LgRF^38,43^. S_ON-OFF_ might correspond to G14 ON-OFF (both predominantly prefer either one of the cardinal orientations, and the response of both is dominated by an ON component)^42^. S_ON-OFF_ V-cells could also correspond to G6 ON-OFF ‘JAM-B’, which exhibits sustained firing and may act as a vertically-preferring OFF OSGC^43,90^, while S_ON-OFF_ H-cells might correspond to the horizontally-preferring OFF OS type^43^. To confirm correspondence between the four OSGC types identified here and those previously documented, it would be beneficial to target cells assigned to each functional type based on Ca^2+^ imaging for detailed morphological and functional analysis, perhaps through whole-cell recording with dye-filling.

### Topographical variation in OSI may represent an evolutionary adaptation for edge detection across the visual field

The strength of OS tuning (quantified by OSI) in the dorsal retina was higher than in the ventral retina, for all four OSGC types. These topographic differences in OS tuning together with known topographical differences in UV cone expression^51–53^, match equivalent differences in the incidence and spectral composition of edges in the natural environment of mice. Utilizing footage from natural habitats of mice, approximating the sensitivity spectra of the mouse UV and M (green) cone opsins^91^, we found that the incidence of edges was higher in the lower visual field in the green channel, but higher in the upper visual field in the UV channel. This coincides with the higher OSI observed in the dorsal retina, along the higher expression of UV cone opsin reported in the ventral retina^51–53^, suggesting that retinal topographic variation in OSI and UV cone opsin may represent an adaptation for the detection of edges in the natural habitat of the mouse.

Additionally, the two ON-OFF types exhibited a gradient of OSI, being highest in the dorsal retina and lowest in the ventro-temporal retina. This OSI gradient might coincide with equivalent differences in the incidence of ipsilaterally-projecting RGCs. While RGCs across the whole retina project contralaterally, ipsilaterally-projecting RGCs are confined to the ventro-temporal retina^92,93^. Therefore, our results suggest that ipsilaterally-projecting OSGCs may exhibit lower OS tuning. However, as the projection patterns of OSGC are still being uncovered ^33^, this possibility is yet to be tested. Also awaiting exploration are the mechainsms underlying the topographical differences in OS tuning, which may include differences in the speed-tuning of OSGCs.

## Methods

### Animals

26 C57BL/6J (Jackson Laboratory) mice (2-2.5-months old, Both sexes). No statistical methods were used to predetermine the sample size. Animals were kept under a 24-hour light cycle (T24) and given water and food ad libitum. Imaging experiments were conducted during the animal’s subjective day. All procedures were in accordance with National Institutes of Health guidelines and approved by the Institutional Animal Care and Use Committee at Brown University.

### Intravitreal injections

C57BL/6J mice were anesthetized with isoflurane (3% in oxygen) using a Midmark Matrx VIP 3000 calibrated vaporizer. The vitreous chamber of their right eye was then injected with 1.5 –2 μl of ∼3 x 10^13^ units/ml of viral vector (AAV2/1.hSynapsin. GCaMP6f; Vector Core, UPenn) through a glass pipette using a General Valve Corporation Picospritzer III microinjector to induce expression of calcium indicator GCaMP6f. The glass pipette was used for puncturing and entering the ora serrata in the ventro-temporal side of the eye.

### Retinal dissections

Two-three weeks following viral injection, animals were sacrificed, and the right eye removed and immersed in an oxygenated Sigma-Aldrich Ames’ medium (95% O_2_, 5% CO_2_) supplemented with 23 mM NaHCO_3_ and 10 mM D-glucose. Then a small incision was made at the ora serrata and the eye cut along it to remove the cornea, the lens and the vitreous humor. To determine the eye’s orientation, the eyecup was cut at four different locations, two cuts at each of the insertions of the lateral and medial recti, and two asymmetrical cuts at the ventral and dorso-temporal parts. The retina was then separated from the eyecup, placed flat on a cell culture insert ^94^ using gentle suction and secured on a custom-build chamber on the microscope stage. While in the chamber, the retina was continuously superfused with Ames’ medium (32-34°C). To measure the arc length from the ora serrata to the optic disk, the left eye was also removed and photographed, for mapping flatmounted retina data into spherical coordinates ^44^. Retinas were used for up to 8 hours. All retinal dissections and experimentation were conducted under dim red light or darkness to minimize unwarranted photoreceptors activation.

### Immunohistochemistry

After experimentation, retinas were fixed in 4% paraformaldehyde for 30 minutes (20°C). To identify ganglion cells, 6 retinas were counterstained with guinea pig anti-RBPMS (RNA-binding protein with multiple splicing; 1832-RBPMS, PhosphoSolutions), a pan-ganglion-cell marker ^95^. Others were stained with chicken anti-GFP (Abcam), to augment the fluorescence GCaMP6f and align cells between experimentation imaging and multiphoton microscope imaging after fixation, for which retinas were oriented under the microscope to replicate experimentation orientation.

### Two-photon functional imaging

Calcium imaging was performed using an Olympus FV1200MPE BASIC (BX-61WI) microscope fitted with an Olympus XLPL25XWMP 25X, 1.05 NA water-immersion objective and a Spectra-Physics Mai Tai DeepSee HP laser line tuned to 910 nm. A 570nm dichroic mirror and a FF03-525/50-32 Semrock bandpass filter were installed before the microscope’s GaAsP photomultiplier tubes (PMTs) to filter out light at a wavelength lower than 525 nm or higher than 575 nm. The imaging system was controlled through FluoView, and images acquired at a zoom of 2 and a scanning frequency of 15 Hz such that each pixel in the image is equivalent to 1 μm on the surface of the retina.

### Visual stimulation

Visual stimuli were generated using custom MATLAB scripts created with the Psychophysics toolbox^96^. Stimuli were projected onto the retina using a DLP AX325AA HP Notebook Projection Companion fitted with a 385 nm UV light emitting diode (LED) through a FF01-440/SP Semrock short-pass dichroic filter and a T425lpxr Chroma dichroic mirror to filter out light at a wavelength higher than 440 nm or lower than 425 nm. The orientation preference of cells was assessed using sinusoidal grating stimuli drifting in eight directions (spatial frequency=0.132 cycle per degree, Michelson contrast= 0.95, stimulus duration= 3.65 s, inter-stimulus duration= 5 s at uniform mean grating luminance, 45° intervals, drift speed= 4.5° s^−1^, 4 repetitions); the preferred orientation was defined as the orientation orthogonal to the gratings’ movement direction. To determine the response polarity of cells, a bright bar on a dark background moving in eight directions was used (bar width= 1,500 μm, inter-stimulus duration= 5 s, 45° intervals, drifting speed= 300 μm s^−1^, 4 repetition).

The stimulus spectrum was measured using an Ocean Optics USB4000-UV-VIS-ES absolute-irradiance-calibrated spectrometer. From it and using the estimated rod (0.85 μm^2^) and cone (1 μm^2^) collecting areas^96^, we estimated the photoisomerization rates, which were similar to those of rods and cones [∼10^4^ photoisomerizations/s (R*/photoreceptor/s)], independent of their relative expression of S- and M-cone pigments^71^. Stimuli were projected at very high frequency (>2000 Hz); well above critical fusion frequency in mice^97^. Stimulus projection and imaging laser scanning were interleaved such that when the laser was scanning, no stimulus was projected; 50 µs of stimulus were projected in the 185 µs gap between laser scans.

### ROI selection and data analysis

Calcium responses were analyzed using FluoAnalyzer^97^ and custom MATLAB functions. For each field of view (FOV), we manually assigned regions of interest (ROIs), each defining the soma of a single cell. The average intensity of the pixel within each ROI was used as proxy for changes in GCaMP6f fluorescence and therefore also for cell activity^98^. Fluorescence was measured as the difference between the instantaneous fluorescence, *F*, and the mean fluorescence over one second before stimulation onset, *F*_0_, such that:

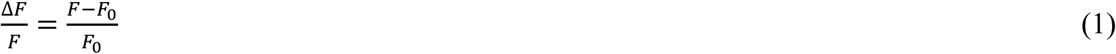

The preferred direction (PD) of a cell was calculated as the angle of the vector sum as follows^28,99^:

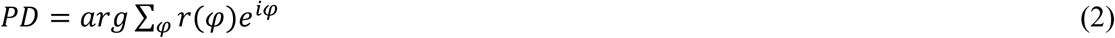

where *r* is the response amplitude to a stimulus moving at a direction *φ* (0°, 45°,…,315°). The direction selectivity index (DSI) of cells which may range between 0 (no direction selectivity) and 1 (highest direction selectivity) was calculated as:

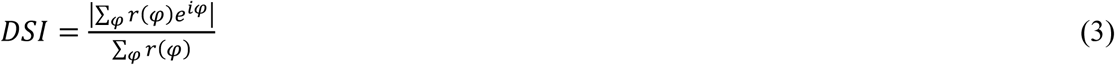

The precise preferred orientation of a cell (*PO*, in degrees), not restricted to the eight probed directions, was estimated by fitting the data to two von Mises distributions (a circular analog to the Gaussian distribution), which take the following form^28,99^:

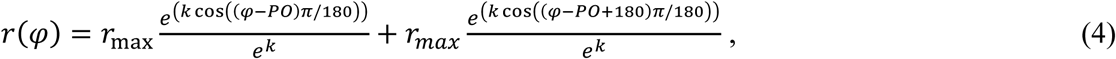

where *r* is the response amplitude to stimuli moving at direction *ϕ* (0°, 45°, …, 315°), *r*_max_ is the maximum response amplitude, and *k* is the width of the orientation tuning curve. This function assumes that the two peak responses are 180 degrees apart, and are of the same amplitude. Both assumptions are reasonable for OS cells. The lower the root mean square error (RMSE) of the fit, the better the two von Mises distributions explain the variation in the data. We then shuffled the responses along the direction (x) axis, fitted the shuffled data to the two von Mises distributions, and by repeating this procedure hundred of times, we calculated the null distribution of RMSE values (100 permutations yield a 1% resolution when calculating the RMSE distribution). Cells for which the RMSE was <5th percentile of the RMSE null distribution were treated as orientation-selective.

The orientation selectivity index (OSI), which similarly to DSI ranges between 0 and 1, was calculated as^28^:

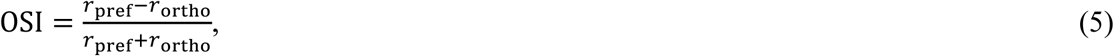

where *r*_pref_ is the maximum response amplitude obtained among the 8 probed directions, and *r*_ortho_ is the mean of responses at the two directions spanning the orthogonal to the direction at which *r*_pref_ was attained. In the above, *r* represents average calcium response from the stimulus onset to 2 sec following stimulus termination, to capture the OFF responses and to account for the slow decay of calcium responses^100^.

### Unsupervised clustering and the diversity of OSGCs

To assess how many subtypes might be differentiable in our sample based on calcium-response kinetics, we performed a principal component analysis (PCA) on the responses of all cells to bar motion in each cell’s optimal tested direction, followed by unsupervised clustering (Gaussian Mixture Models) on the scores of the principal components (PCs) that accounted for over 99% of the variance. For clustering, we used the full covariance matrix, regularized by the addition of a constant (0.001) to guarantee that the estimated covariance matrix is positive. To avoid local minima, we repeated the algorithm using 50 different sets of initial values, with the accepted fit being that with the largest log likelihood. The optimal model (having the optimal number of clusters), was estimated based on the Bayesian information criterion (BIC)^101^.

### Registration of calcium imaging data

To avoid deviations in the preferred orientations of the OSGCs relative to retina’s orientation, orientations of all retinas were corrected by rotating them based on the displacement of each retina (*n*=26) from a reference map^101^.

### Assessing the alignment between OSGC preferences and the longitudinal, latitudinal, and hybrid model families

We assessed the alignment between OSGC preferences and three families of models: longitudinal, latitudinal, and hybrid. Each family consisted of 684 models, also called axes, defined by spherical angles along direction (0–350°) and eccentricity (0–180°) at 10° intervals. Each axis represents a population of modelled cells with locations matching those of OSGCs and preferred orientations aligned with specific field line patterns – either lines of longitude or latitude. Due to technical issues, two instead of one modelled cell were generated for each real cell. However, since this doubling was applied to all real cells, it did not affect any of the analyses. To quantify the degree of alignment between the preferences of OSGCs and the modelled cells corresponding to a specific axis, we introduced a concordance index defined as the percentage of OSGCs with orientation preferences within ±10° of their corresponding modelled cell. Repeating such template-matching process across all axes of the model family, we created a tuning map displaying concordance as a function of the model axis. Hotspots on the tuning map represent the axes that formed modelled cells that most closely aligned with OSGC preferences, reflecting higher concordance.

### Estimating the relative abundance of single subtypes in samples of OSGCs

For each OSGC type independently, we generated a modelled tuning map comprising two OS subtypes (H- and V-cells), each of which aligned its OS preferences everywhere with the field lines produced by longitudinal (translatory-like) geometry, latitudinal (rotatory-like) geometry, or hybrid geometry, along a single, empirically determined best axis. For hybrid geometry only one subtype was used since we assume that the subtypes’ OS preferences are always orthogonal and therefore the field lines that they align with originate from or circulate around the same axis. Coordinates of these two best axes were derived from the two local maxima (hotspots) in the tuning maps. We asked what weighting of these subtypes could best reproduce the longitudinal-geometry or latitudinal-geometry tuning map for all OSGCs. Formally, the best fitting tuning map for modeled cells (*M*_f_) was derived from the weighted sum of two (i=1,2) single-subtype maps *M*_i_ (Figure 3e-h):

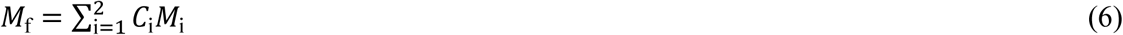

where *C*_i_ denotes the weighting allocated to individual subtype maps. We used least-squares fitting (regress function in Matlab) and the multiple correlation coefficient (*R*^2^) for regression models without a constant term to assess the goodness of fit ^102^. To enable comparison between the hybrid model and the longitudinal or latitudinal models, that differed in the number of regressors, the *R*^2^ was adjusted as:

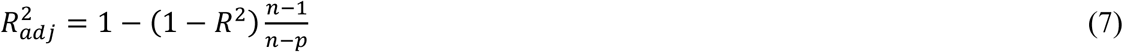

where *n* is the sample size (number of cells tested) and *p* is the number of regressors (number of axes). Two-tailed 95% confidence intervals (CI) for the adjusted *R*^2^ values were calculated using bootstrapping. Specifically, for each OSGC type, we resampled the actual OSGCs with replacement, generated a tuning map, fitted that map with the two individually-weighted single-subtype maps comprising the model, and determined the coefficient of determination (*R*^2^) that represents the model’s fit to the data. We repeated this procedure 1000 times and calculated the CI of *R*^2^. CI were determined in the same way for the longitudinal, latitudinal, and hybrid geometry models. In the case of the hybrid geometry model, the tuning map was fitted by only one weighted single-subtype map that comprised the model.

### Retinal coordinates

The retinocentric coordinate system we used was monocular, had its origin at the nodal point of the eye, and specified retinal or visual-field location by two coordinates: eccentricity and direction. Eccentricity was defined as the visual angle between the point of interest and the optic axis (the projection of optic disk). Eccentricity is 0° at the blind spot and 90° at the margin of the visual hemifield, corresponding to the retinal margin. Direction refers to the orientation of displacement of the point of interest from the optic axis, where nasal=0°, dorsal=90°, temporal=180° and ventral=270°.

### Estimation of the variability in the preferred orientation of OSGCs

Variability in the preferred orientation of OSGCs was estimated based on a subset of 67 OSGCs belonging to all four types. Specifically, for each OSGC, we used bootstrapping to sample with replacement the four repetitions (each comprising the responses to 8 directions), calculated the mean response over the 8 tested directions, fitted it to a double Gaussian function, and determined the OSGC’s preferred orientation. After repeating this procedure 100 times for each OSGC, we calculated the standard deviation of the preferred orientation of the OSGC in question. Lastly, averaging the SD of preferred orientations across all the 67 cells yielded the value 4.4567°, taken as an approximation of the variability in the preferred direction of OSGCs.

### Estimation of dendritic tree dimensions

Dendritic field (circle-equivalent) diameter of OFF-vOS and OFF-hOS cells was reported to equal 239 μm and 224 μm, respectively ^43^. Dendritic field (circle-equivalent) of the bistratified ON-OS cell was reported to equal 195 μm (OFF arbor) and 252 μm (ON arbor) for vertical-preferring cells, and 226 μm (OFF arbor) and 339 μm (ON arbor) for horizontal-preferring cells ^38^. Thus, the dendritic field diameter of OSGCs ranges 195 - 339 μm, across OFF-OS and ON-OS types^38,43^, corresponding to visual angles of 5.9-10.3° (estimated according to image magnification of 0.033 mm/degree at 60 days old mouse^103^). The timepoints of the bar’s entry and exit of the receptive field (RF) were estimated based the bar’s speed and the cell’s position within the imaged field, assuming a 300-μm RF diameter.

### Detection of edges in the natural visual environment of mice

See Qiu et al.^70^ for details on camera design, video recordings, temporal and spatial alignment, and spectral and intensity calibration for the dataset of footage from natural habitats of mice. Two crops (128 x 128 pixels, equivalent to a solid angle of 53°), from the image’s upper and lower halves were extracted from each image. The images brightness varied across times of day, weather, and scene content. Therefore, image crops were divided, already in the original study, into low, median, and high ‘intensity classes’^70^. To avoid potential confounding of the detection of edges by overexposure, we analyzed 1000 randomly-sampled crops comprising the ‘low’ and ‘intermediate’ intensity groups. To detect edges we used the Canny Edge Detection^78^ as implemented for Python in the OpenCV library ^79^ version 4.7.0.72. The edge detector requires specifying two hysteresis threshold parameters, set to 100 and 200. We computed edges for the UV and green channels separately.

## Supporting information

Supplementary Equations

## Acknowledgements

We thank Thomas Euler who made available previsouly-published footage from natural habitats of mice. We are thankful for our colleagues who provided invaluable theoretical, data-analytic and technical advice, Alex Binshtok, Ariel Gilad, Dan Rokni, Yoram Ben Shaul and Eli Shmueli. This project was supported by the Brain & behavior Research Foundation (BBRF) grant (#30072) and the Israel Science Foundation (ISF) grant (#1134/21), awarded to S.S.

## Author Contributions

S.S., D.D.L., and J.A.G. designed the study and developed the theoretical framework. S.S. performed functional imaging experiments and immunostaining and imaging of retinas. D.D.L. and S.S. analyzed all morphological and imaging data. Y.M. developed routines for the detection of edges in the natural visual environment of mice. J.A.G. developed the geometrical decoder and various algoritms used for OSGCs mapping. D.D.L, J.A.G., and S.S. wrote the paper.

## Supplemental information

### Supplementary Figures

Figures S1–S10

### Supplementary Videos

**Video S1.** The probability density representing the distribution of decoded edge orientations (*υ*), as a function of the angle between the mean preferred orientations of the sensors (*ϕ*), and the angles between the real-world edge and the preferred orientation of the sensors (*θ* and *ϕ* − *θ*); related to Figure 6.

**Video S2.** Footage of a mouse visual habitat, in the UV and Green channels, individually and merged, and with the detected edges, for the upper and lower visual space; related to Figure 7.

