## Supplementary Figures for "Spherical Code of Retinal Orientation-Selectivity Enables Decoding in Ensembled and Retinotopic Operation"

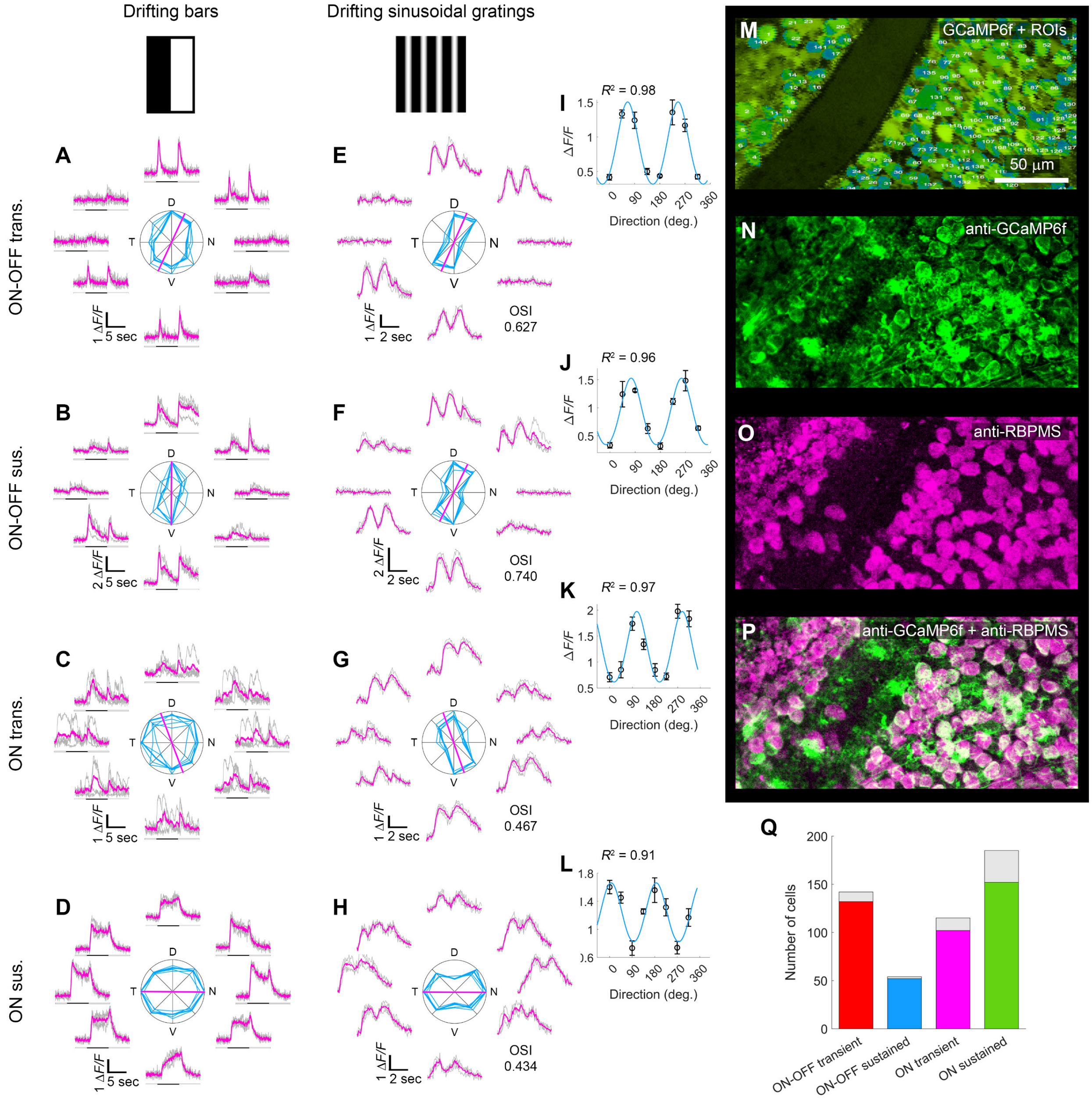


**Supplementary Figure 1. Calcium transients of representative OS cells, and correspondence between calcium-imaged cells and a pan-ganglion cell immunoreactivity**

**A-L**. Calcium transients of representative OS cells of the four functional types in response to bright bars (**A-D**) or sinusoidal contrast gratings (**E-H**) drifting in eight directions at 45° intervals. Preferred orientation and orientation selectivity index (OSI) were determined from grating responses, cell class ($T_{ON\text{-}OFF}$, $S_{ON\text{-}OFF}$, $T_{ON}$, $S_{ON}$) from the bar responses. Traces plot somatic GCaMP6f $Ca^{2+}$ signal over time; magenta shows mean; gray, single trials. Black marker indicates when stimulus bar was within the cell's receptive field. Polar plots show response amplitude (normalized to maximum) for each direction (bold curves, mean of four repetitions; thin, single trials). Magenta vector shows mean preferred orientation (N, nasal; D, dorsal; T, temporal; V, ventral). **I-L.** fit of two Gaussians to the mean response amplitude to drifting grating, with the coefficient of determination, $R^2$ indicated. **M-Q.** Correlating $Ca^{2+}$ imaging and post hoc immunofluorescence in a single imaged field. **M.** A two-photon image of GCaMP6f-expressing cells in live retina, *in vitro*. Numbers indicate regions of interest (ROIs) for assessing somatic OS responses. Cell-free zones are blood vessels. **N–O.** A confocal image of the same field after fixation, immunolabelling, and alignment with the live image (**M**). **N.** Anti-GFP immunofluorescence, to enhance GCaMP6f signal which faded after fixation. **O.** Immunolabelling for RBPMS, a pan-ganglion cell marker. **P.** A merged image combining GCaMP6f (anti-GFP) signal with RBPMS. **Q.** Number of OSGCs per type, immuno-positive (colored bar) and -negative (gray bar) for RBPMS.

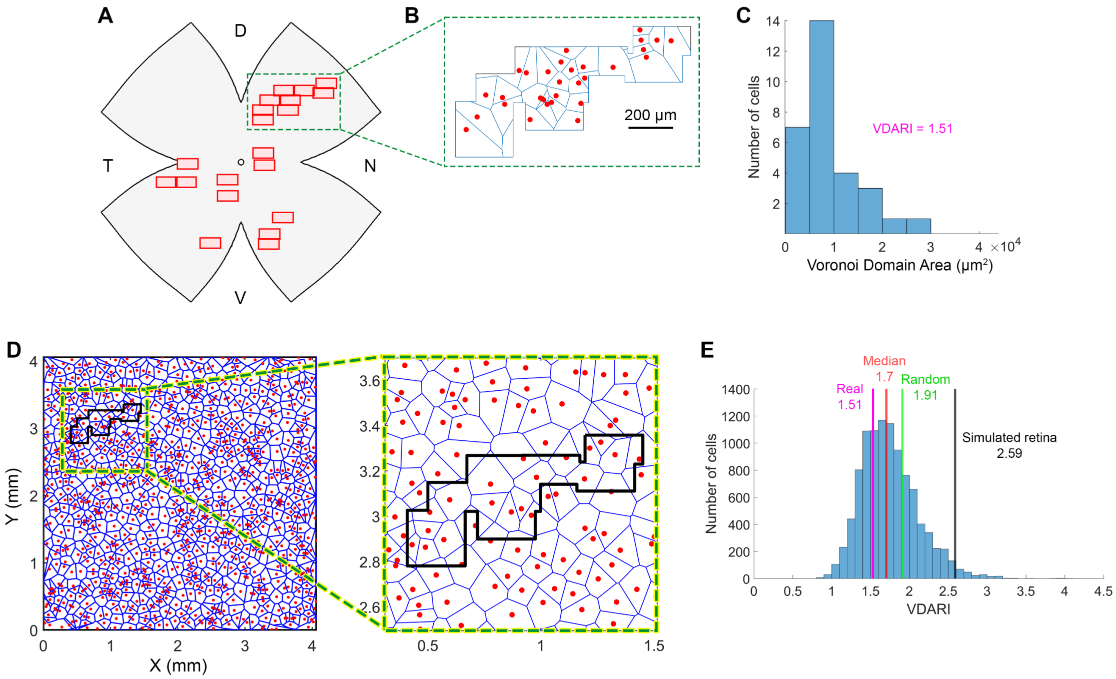


**Supplementary Figure 2. Assessment of retinal mosaics**

Retinal cell mosaics are spatially organized rather than random, optimizing sensory input processing[1,2]. Their organization is commonly assessed using Voronoi domain areas, where each cell's area consists of all points closer to it than any other. The boundaries between these areas form polygons[3]. The Voronoi Domain Area Regularity Index (VDARI) quantifies regularity by dividing the average of all Voronoi domain areas by the standard deviation of areas. Higher VDARI values reflect greater regularity. A cutoff value of VDARI = 1.91 distinguishes random from uniform distributions: values below 1.91 indicate clustering, while values above 1.91 indicate uniform spacing and mosaic formation[4]. **A.** To assess the feasibility of accurately calculating the VDARI based on our non-contiguous data, as the best-case scenario, we selected the retina (out of 28) with the greatest number of adjacent imaged fields of view (FOVs). This retina included 20 FOVs, 9 of which were adjacent, forming a polygonal area of 0.28 $mm^2$. **B.** We analyzed cells of the $T_{ON\text{-}OFF}$ type which was the most abundant OSGC type (30 cells) within the polygon. **C.** The mean VDARI of these $T_{ON\text{-}OFF}$ cells was calculated by be 1.51, below the random distribution threshold (1.91), indicating a clustered cell distribution. **D.** To determine the likelihood of obtaining a VDARI corresponding to a uniform distribution from non-adjacent FOV data, we simulated a retina with cells uniformly-distributed, sampled it 10,000 times using a moving polygon matching the real envelope polygon's size, and calculated VDARI for each iteration. The simulated retina included 1,000 cells in a 16.5 $mm^2$ square, equivalent to the C57BL/6 mouse retina surface area[5]. By adjusting intercellular distances, the simulated VDARI was set to 2.59 (black line), similar to that observed in JAM-B RGCs (2.7), which may be analogous to $T_{ON\text{-}OFF}$ OSGCs[6]. **E.** Distribution of 10,000 simulated VDARI values. The median of simulated VDARI values (1.7, red line) and 71.36% of the simulated VDARI values were below the random distribution threshold (1.91, green line). This indicates that, even with a uniform cell distribution, the small size and irregular shape of the envelope polygon often produced VDARI values corresponding to clustered distributions. Thus, although the real VDARI for $T_{ON\text{-}OFF}$ cells in the example retina (1.51, magenta line) was found smaller than the random distribution threshold (1.91), it remains inconclusive whether these cells as well as cells of different types from other retinas (where the numbers of adjacent FOVs are even lower) are uniformly spaced.

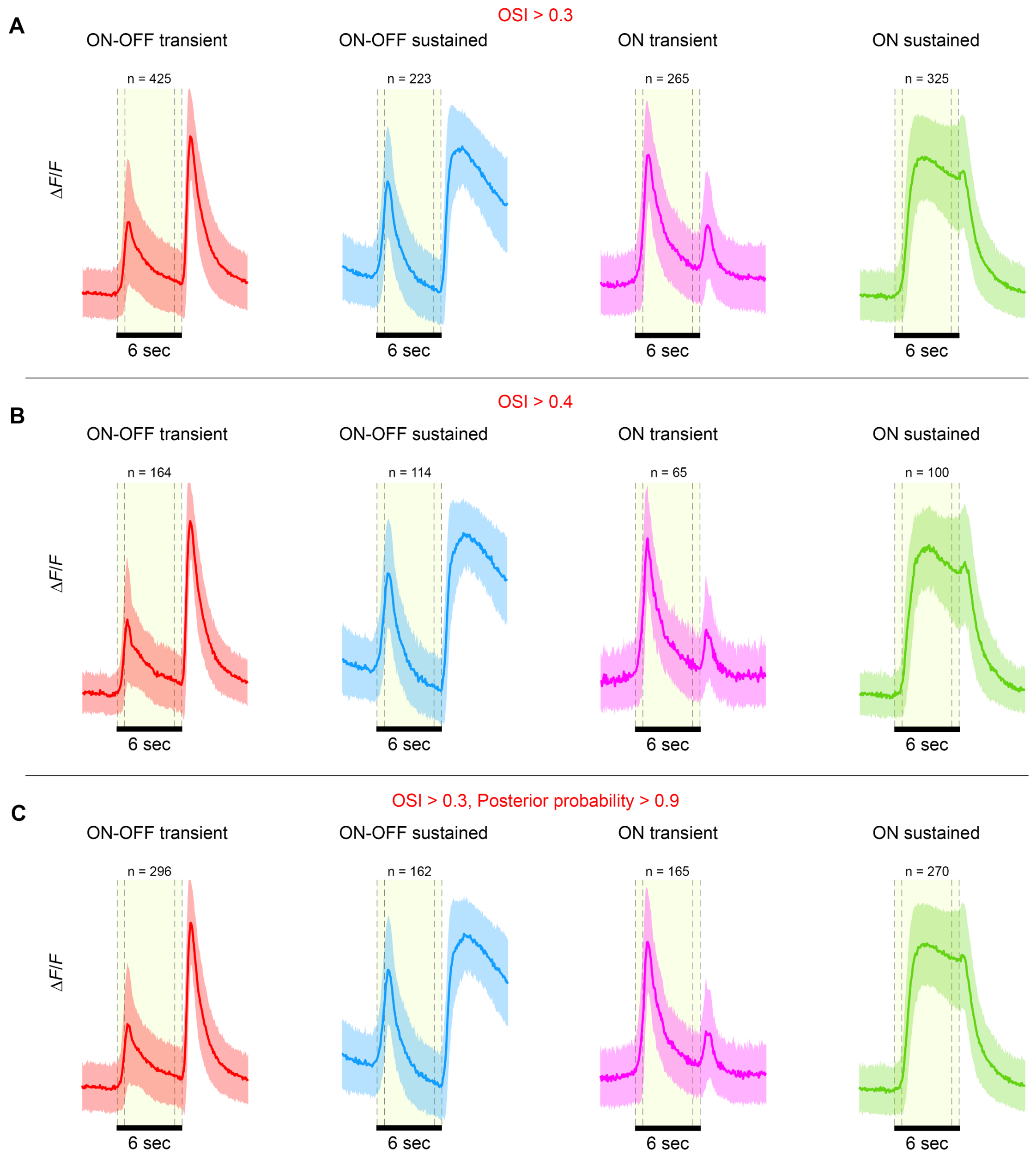


**Supplementary Figure 3. Kinetics of the four OSGC types is similar across OSI and posterior probability criteria.**

Response to bar motion only for cells with OSI>0.3 (**A**), only for cells with OSI>0.4 (**B**), and only for cells with OSI>0.3 and a posterior probability>0.9 to belong to each of the types (as opposed to the default posterior probability>0.5 criterion) (**C**). **All panels.** Calcium responses [$\Delta F/F$, mean (line)±s.d. (shaded area)] evoked by a light bar moving in the cell's preferred orientation, for each of the four identified OSGC types: $T_{ON\text{-}OFF}$ (red), $S_{ON\text{-}OFF}$ (blue), $T_{ON}$ (magenta), and $S_{ON}$ (green). Shaded yellow area and dashed black lines mark the approximate time when the stimulus bar was within the cell's receptive field.

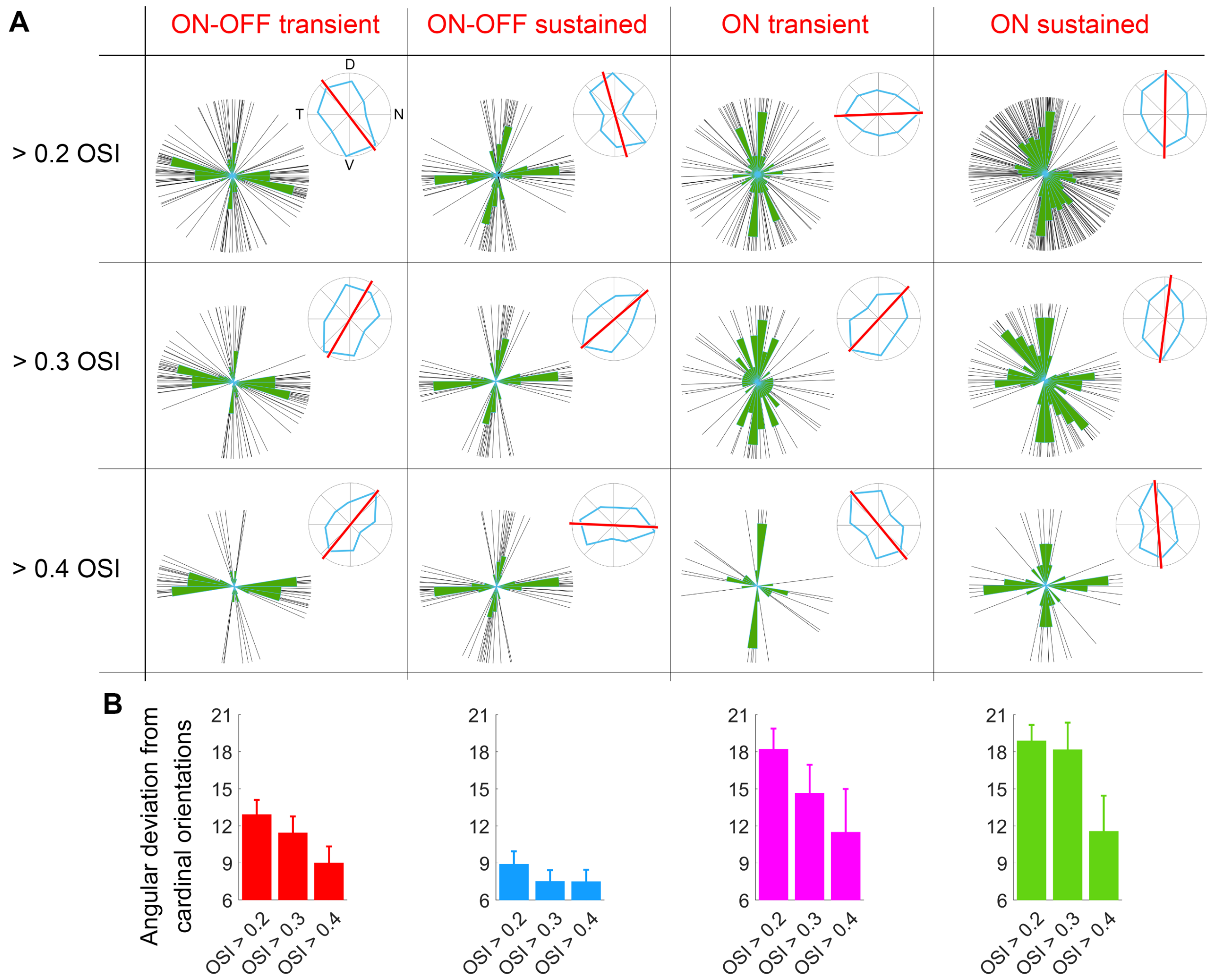


**Supplementary Figure 4. Variation in orientation selectivity index (OSI) across central-retina OSGCs**

OSI varies between studies, stimuli, and recording methods[7-11], and thus the adoption of previously-employed OSI thresholds might be suboptimal for the designation of OSGCs in the current study. To overcome this, for each OSGC type, we evaluated the variation in the mean angular deviation of the preferred orientations of central-retina OSGCs from the cardinal orientations, across 3 OSI threshold values (0.2, 0.3 and 0.4). Based on this procedure, we designated all cells with OSI>0.3 as OSGCs. **A.** OS preferences of individual central-retina OSGCs (sampled region as in Fig. 3C) of the four types, that have survived thresholding at 3 OSI values. Associated polar histograms are overlaid. **Top-right.** Example polar tuning map (blue line) and preferred orientation (red line) of a single OSGC, that was sampled from the entire retina, constructed based on the cell's response to drifting sinusoidal grating stimuli, and survived the OSI threshold in question. **B.** Mean angular deviation of the central-retina OS preferences from cardinal orientations (horizontal and vertical) across OSI threshold values, for each OSGC type. Mean angular deviation generally decreases with increasing OSI threshold.

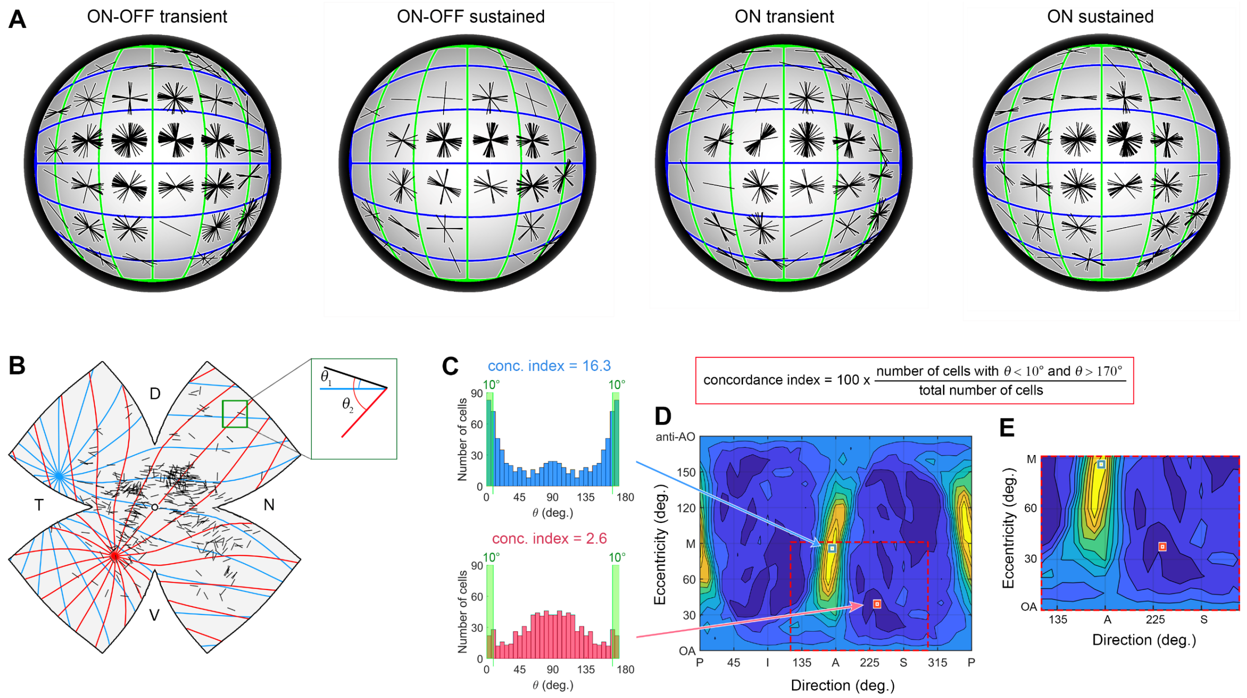


**Supplementary Figure 5. Development of a tuning model, and topographic dependence spherical retina**

**A.** Topographic dependence of orientation preferences, displayed on a projected spherical retina, for each of the four OSGC types. For reference, green and blue lines represent orthogonal longitudinal field lines. **B-E.** Generation of tuning maps. **B**, OS preferences of all imaged cells of the $T_{ON\text{-}OFF}$ type mapped onto a standard flattened retina. Each black line marks one cell's location and preferred orientation. How well do these OS preferences align with the local retinal optic pattern produced by three families of geometric models (longitudinal, latitudinal, hybrid)? In this example, two axes defining two longitudinal fields are shown (purple and yellow lines); many more were tested [684 axes; 10° intervals of spherical angle along the direction (0-350°) and eccentricity (0-170°) coordinates]. For each tested longitudinal model, we measured the angle $\theta$ between each cell's preferred orientation and the local orientation of the longitudinal optic pattern (inset: $\theta_1$ for the purple field, $\theta_2$ for the yellow one). N, nasal; D, dorsal; T, temporal; V, ventral. **C.** Distributions of angles $\theta_1$ (top) and $\theta_2$ (bottom) among all cells in (**B**). Concordance index constitutes the percentage of cells with OS preferences differing less than 10° or more that 170° from the local orientation in a specific longitudinal field (green rectangles). Alignment was much greater with 'purple' than 'yellow' longitudinal field (concordance index=16.3 and 2.6, respectively). **D.** Tuning map displaying concordance index as a function of the axis of longitudinal geometry model. Location of data for 'purple' and 'yellow' longitudinal fields are indicated. Direction represents polar direction in the visual field, e.g., S, superior (corresponding to ventral retina); A, anterior (temporal retina); I, inferior (dorsal retina); and P, posterior (nasal retina). Eccentricity is 0° at the projection of the optic disk onto central visual field (OA) and 90° at the margin of the visual hemifield (M), corresponding to the retinal margin. Eccentricities of 90°–180° correspond to spatial locations outside the visual hemifield. Thus, the lower half of the tuning map represents the visual hemifield, while the upper half represents the 'blind hemifield' of extrapersonal space, not seen by the retina. **E.** The full-range tuning map was restricted to 120-300° along the direction coordinate and to 0-90° along the eccentricity coordinate to include only one pair of unique hotspots (best axes).

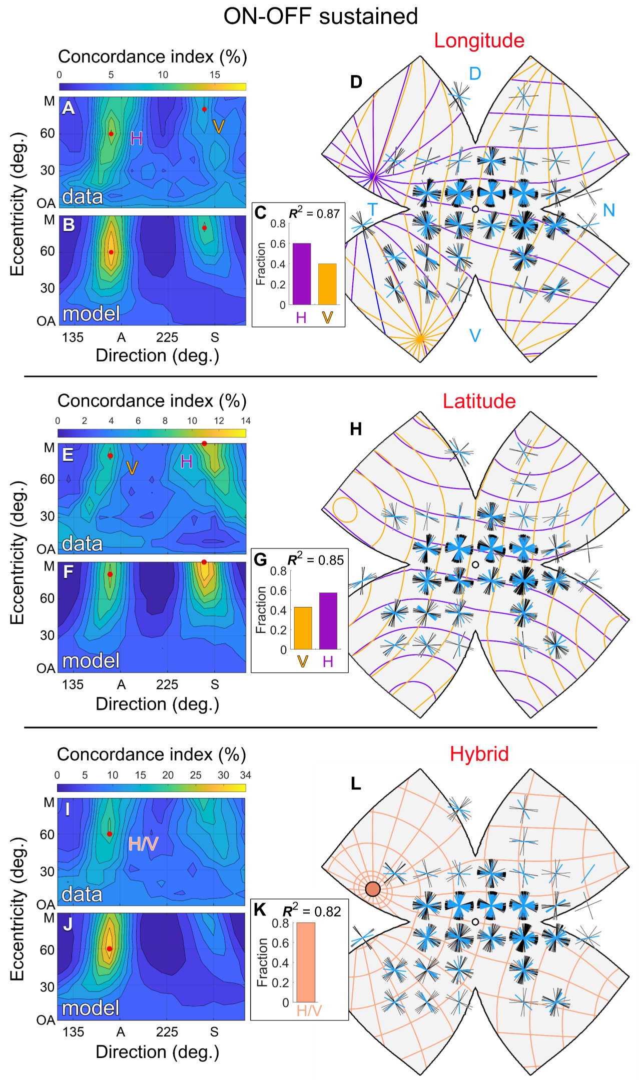


**Supplementary Figure 6. OS preferences of $S_{ON-OFF}$ OSGCs align best with longitudinal geometry.**

**A**. Concordance index based on the orientation preferences of $S_{ON-OFF}$ OSGCs (n = 223) as a function of the axis of the longitudinal geometry model. **B**. Concordance index tuning map of OS preferences of two subtype ensembles (H-cells, V-cells). **C**. Relative weighting of the different subtypes (H-cells, V-cells) comprising the model in (**B**). **D**. Flattened standardized retina overlaid with a grid of local orientation preference polar plots for the modelled cells (black) and for the real cells (blue). The best fitting longitudinal axes are represented by purple and yellow field lines. **E-H**. Same as (**A-D**) for the latitudinal geometry model. **I-L**. Same as (**A-D**) for the hybrid geometry model.

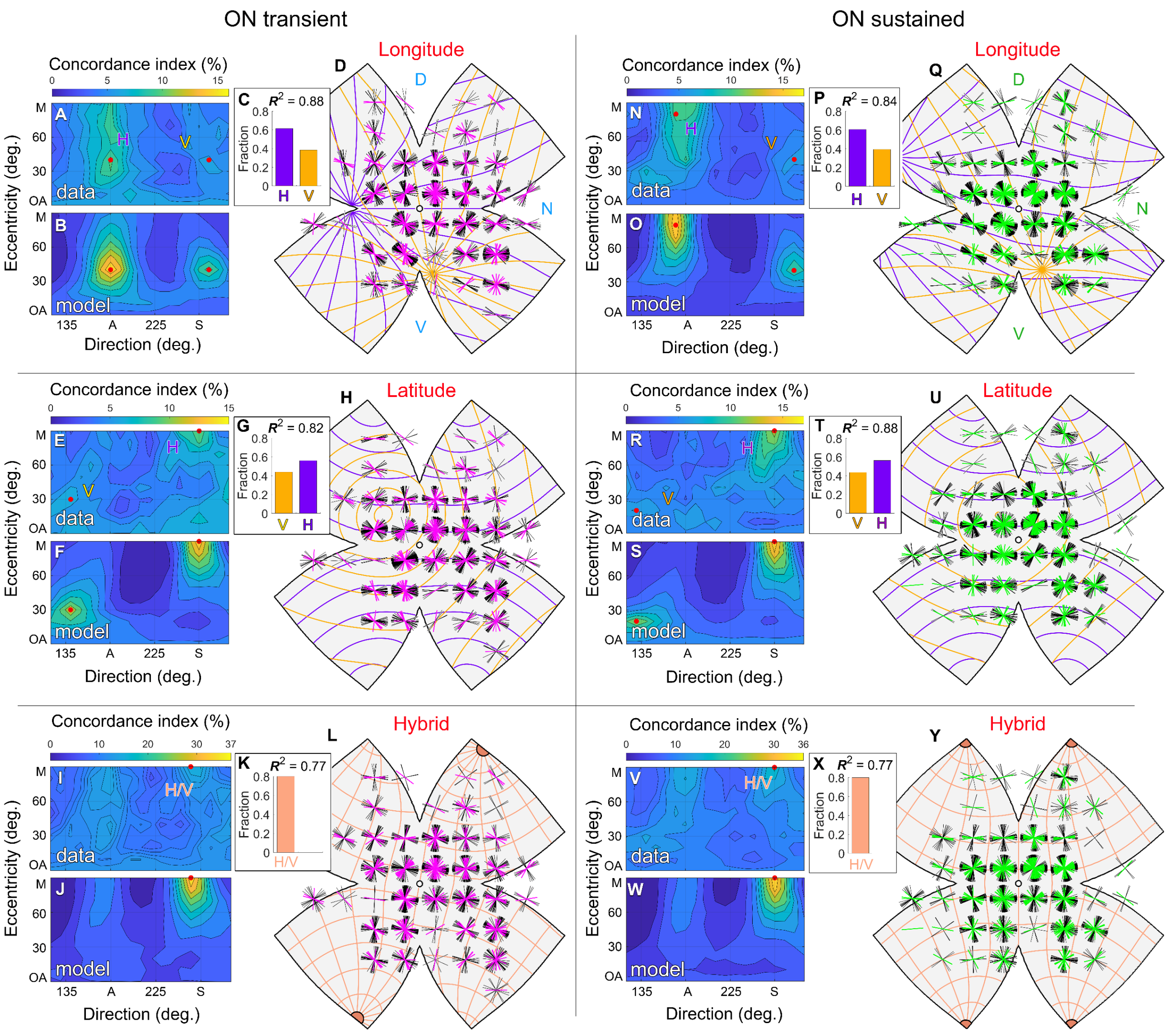


**Supplementary Figure 7. OS preferences of $T_{ON}$ OSGCs align best with longitudinal geometry, while those of $S_{ON}$ OSGCs align best with latitudinal geometry.**

**A,N**. Concordance index based on orientation preferences of $T_{ON}$ (n = 265, **A**) and $S_{ON}$ (n = 325, **N**) OSGCs, as a function of the axis of the longitudinal geometry model. **B,O**. Concordance index tuning map of orientation preferences of modelled $T_{ON}$ and $S_{ON}$ types as a function of the axis of the longitudinal geometry field. **C,P.** Relative weighting of the two subtype ensembles (H-cells and V-cells) comprising the model in (**B,O**). **D,Q**. Flattened standardized retina overlaid with a grid of local orientation preference polar plots for the modelled cells (black) and for the real cells (purple, $T_{ON}$; green, $S_{ON}$). The best fitting longitudinal axes are represented by purple and yellow field lines. **E-H, R-U**. Same as (**A-D, N-Q**) for the latitudinal geometry model. **I-L, V-Y.** Same as (**A-D, N-Q**) for the hybrid geometry model.

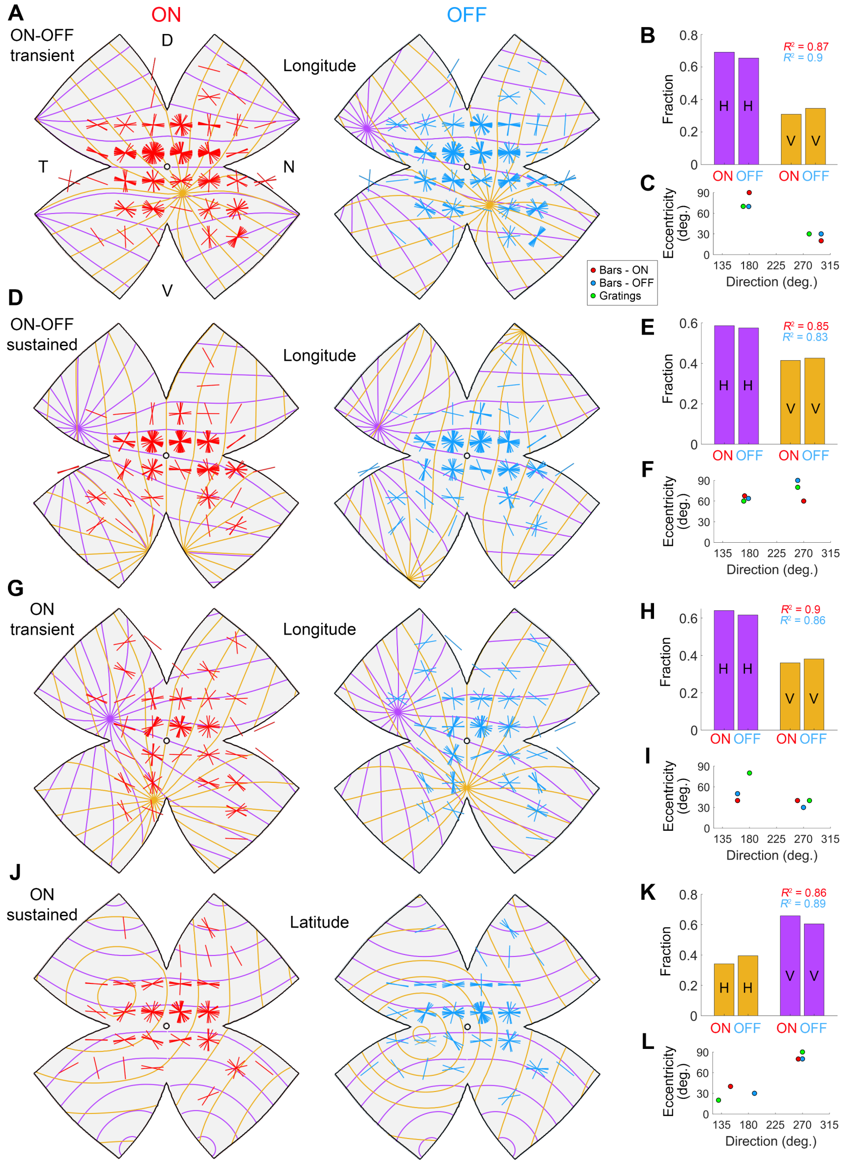


**Supplementary Figure 8. Comparison of orientation preferences determined based on bar and grating motion.**

**A**. Standardized retinas with a grid of local orientation preference polar plots for $T_{ON\text{-}OFF}$ cells, based on their ON (red, left) and OFF (blue, right) components in response to bar motion. The ON and OFF components were calculated as the average response over two 4.05-sec windows encompassing the rise and peak of the ON and OFF responses, and only cells with OSI>0.1 were included in further analyses. The best fitting longitudinal axes are represented by purple and yellow field lines. **B.** Relative weighting of the different subtypes (H-cells, V-cells) comprising the best fitting model, for the ON and OFF response components. **C.** Retinal coordinates of best-aligned axes (hotspots) for $T_{ON\text{-}OFF}$ as determined based on the ON and OFF components in response to bar motion, and in response to grating motion (results for grating motion are similar to those presented in Fig. 5A). The models' goodness of fit ($R^2$) is indicated. **D-F.** Same as (**A-C**) for alignment of $S_{ON\text{-}OFF}$ cells with the longitudinal geometry model. **G-I.** Same as (**A-C**) for alignment of $T_{ON}$ cells with the longitudinal geometry model. **J-L.** Same as (**A-C**) for alignment of $S_{ON}$ cells with the latitudinal geometry model.

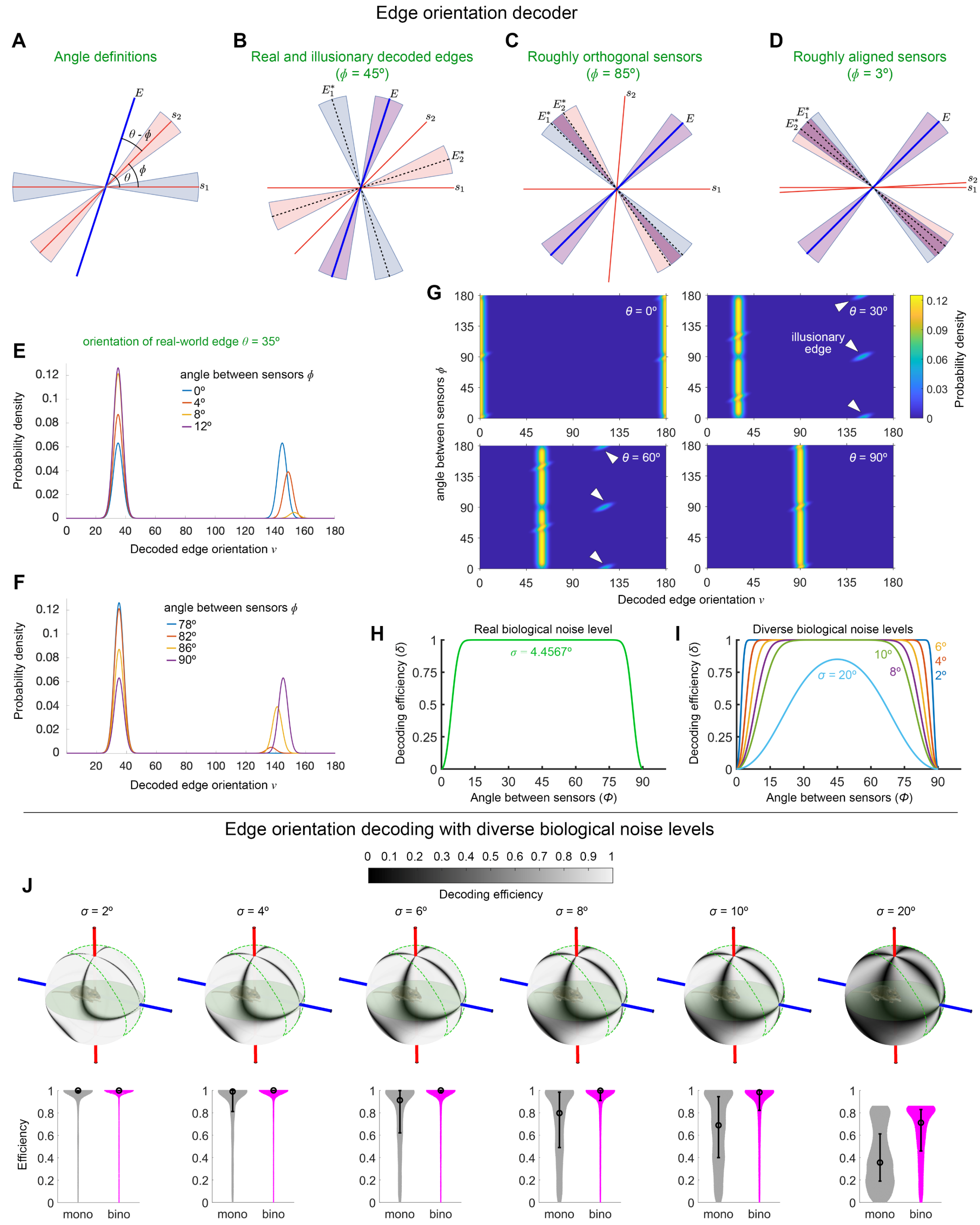


**Supplementary Figure 9. Edge orientation decoder**

**A.** Schematic for encoding of the orientation of a real-world edge into responses of two OS sensors

($s_1$ and $s_2$). Blue and pink shaded regions around the sensors illustrate the uncertainty in the preferred orientation of the sensors, having an azimuth width of 4 standard deviations (4σ). The random variables $\theta$ and $\phi$ correspond to the orientation of the edge measured with respect to the preferred orientation of $s_1$ and the angle between the preferred orientations of $s_1$ and $s_2$, respectively. **B-D.** Schematic for decoding of edge orientation from the responses of the sensors ($s_1$ and $s_2$) for varying angles between the two sensors $\phi$=45° (**B**), 85° (**C**), and 3° (**D**). The blue and pink shaded regions illustrate the uncertainty in the decoded orientation of the real edges $E$ and illusionary edges $E^*$, having azimuth widths of 4σ. The purple shaded regions, overlaid on top of the blue and pink regions, correspond to the overlap of the support of each density for the decoded ($\theta$). Specifically, the purple region illustrates the most probable orientations decoded based on the responses of the two sensors. **E.** Example probability densities of orientations decoded based on the responses of two sensors that are roughly aligned (angles between sensors $\phi$=0°, 4°, 8°, and 12°; orientation of a real-world edge was 35°). With $\phi$=0° (blue), the probabilities of decoding the real and illusionary edges are equal; with increasing $\phi$, the probability of decoding the real edge increases, and that of the illusionary edge decreases, with the probability of decoding the illusionary edge approaches 0 at $\phi \approx 12$. **F.** Same as (**E**) but for two sensors that are roughly orthogonal (angles between sensors $\phi$=78°, 82°, 86°, and 90°; orientation of a real-world edge was 35°). With $\phi$=90° (purple), the probabilities of decoding the real and illusionary edges are roughly equal; with decreasing $\phi$, the probability of decoding the real edge increases, and that of the illusionary edge decreases, with the probability of decoding the real edge approaches maximum at $\phi \approx 78$. **G.** Example snapshots from Video S1 showing the probability density as a function of the angle between sensors ($\phi$) and the decoded edge orientations ($\upsilon$), for real-world edge orientations $\theta$=0°, 30°, 60°, and 90° (orientation of the real-world edge relative to the preferred orientation of sensor 1). Arrowheads mark the probability densities and the associated decoded orientations of illusionary edges. **H,I.** Efficiency of edge orientation decoding ($\delta$) as a function of the angle between the sensors ($\phi$), for the empirically-determined level of biological noise ($\sigma$) (**H**) and a series of theoretical biological noise levels ($\sigma$=2°, 4°, 6°, 8°, 10°, and 20°) (**I**). **J.** Decoding efficiency of ${}_{long}$H/${}_{long}$V-cells of the right eye while accounting for a series of theoretical biological noise levels ($\sigma$=2°, 4°, 6°, 8°, 10°, and 20°). Decoding efficiency in the binocular zone was significantly higher than in the monocular zone (permutation t-test, $p<0.0001$, $n_{mono}$=46400, $n_{bino}$=18941, across all tested biological noise levels).

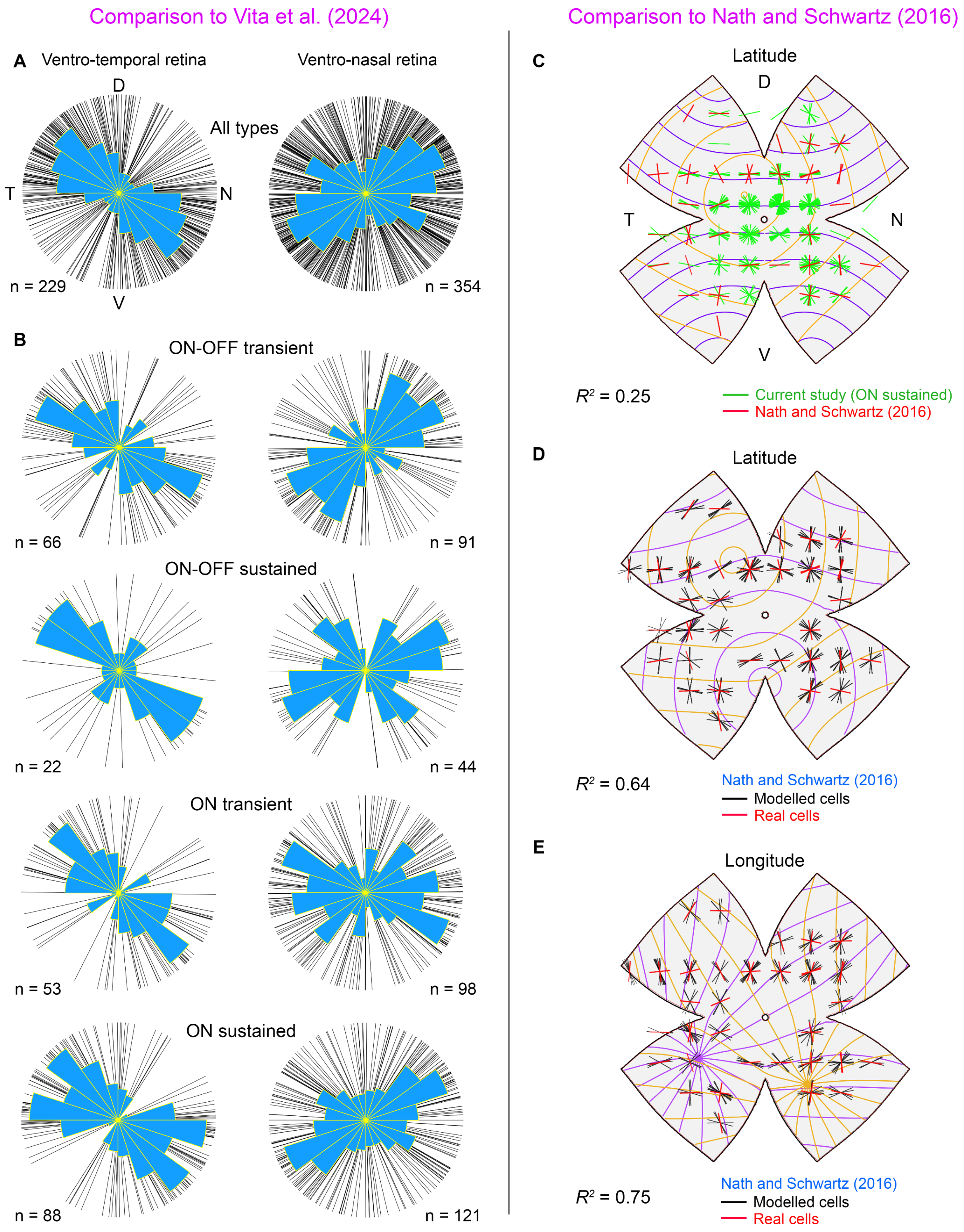


**Supplementary Figure 10. Comparison to previous studies**
**A,B.** Comparison of prefereed orientation in the ventro-temporal and ventro-nasal retina, between the current study and Vita et al. (2024), across all OSGCs collectively (**A**) and within each of the four identified OSGC subtypes (**B**). In general, here and as reported in the previous study, OSGCs in the ventro-nasal retina preferred orientations along the ventro-temporal to dorso-nasal axis, whereas OSGCs

in the ventro-temporal retina preferred orientations along the ventro-nasal to dorso-temporal axis. **C-E.** Comparison of preferred orientation across the retina, between the current study and Nath and Schwartz[12]. **C.** Flattened standardized retina overlaid with a grid of local orientation preference polar plots for the ON sustained OSGCs in the current study (green) and for the ON OS cells reported in the previous study (n = 48, red). Preferred orientations of ON OS cells aligned poorly ($R^2$ = 0.25) with the the latitudinal axes (purple and yellow field lines) that we found to fit best to the preferred orientations of ON sustained OSGCs. **D.** Flattened standardized retina overlaid with a grid of local orientation preference polar plots for the real cells in the previous study (red), and for the cells modelled (black) based on the best fitting latitudinal axes (**D**) and longititudinal axes (**E**) (purple and yellow field lines). Preferred orientations of ON OS cells aligned poorly with the the best fit latitudinal axes ($R^2$ = 0.64) and longitudinal axes ($R^2$ = 0.75).
