## Supplementary Equations for "Spherical Code of Retinal Orientation-Selectivity Enables Decoding in Ensembled and Retinotopic Operation"

### Supplementary Equations: Encoding and Decoding of Edge Orientation

In this section, we propose: (1) a procedure for encoding edge orientation by OSGCs whose preferred orientation is subject to biological noise, and (2) a procedure for decoding edge orientation by neurons postsynaptic to OSGCs.

#### Encoding of Edge Orientation

We begin with a setup for the geometry used in the encoding procedure as well as the framework for the probability model we use. Note, we assume that all angles are measured in degrees but, to be consistent with the rules of calculus, we assume all trigonometric functions take in inputs in radians. Since we are concerned with the absolute angular difference between lines, all angles will be measured from $0°$ to $180°$.

Based on the analysis presented in the main text, we assume that the preferred orientation of each cell follows a von Mises distribution with standard deviation $\sigma = 4.4567°$ and at each location in the retina there are two OSGCs $s_1$, $s_2$ with different preferred orientations. Recall, that the probability density for a von Mises distribution is given by

$$f(\theta; \mu, \kappa) = (180° I_0(\kappa))^{-1} \exp\left(\kappa \cos\left(\frac{\pi}{90°}(\theta - \mu)\right)\right), \tag{1}$$

where $\kappa = (90°/(\sigma\pi))^2$ and $I_0$ is the zeroth order modified Bessel function of the first kind [S1]. We choose a von-Mises distribution since, for the values of $\kappa$ we consider, it is an accurate approximation of a wrapped Gaussian distribution with standard deviation $\sigma$ [S2, S3].

Without loss of generality, we assume a coordinate system at each location in the retina in which the mean of the preferred orientation of $s_1$ is aligned with the horizontal axis. We further let $\phi$ denote the angle between the mean preferred orientations of $s_1$ and $s_2$. Consequently, in this coordinate system, the preferred orientations of $s_1$, $s_2$ are independent random variables $\Theta_1, \Theta_2$ with the following von Mises probability densities:

$$\Theta_1 \sim f(\theta_1; 0, \kappa) \text{ and } \Theta_2 \sim f(\theta_2; \phi, \kappa). \tag{2}$$

Furthermore, we assume that a real world edge makes an angle $\theta$ and $\phi - \theta$ with respect to the mean preferred orientations of $s_1$ and $s_2$. Consequently, given the uncertainty in $s_1$ and $s_2$, it follows that the angles the edge makes with each of the two sensors are independent random variables $\Theta$, $\Phi$ with the following probability densities

$$\Theta \sim f(\theta_1; \theta, \kappa) \text{ and } \Theta - \Phi \sim f(\theta_2; \phi - \theta, \kappa). \tag{3}$$

In Supplementary Figure 7A we present an illustration of the geometry used in this model. The uncertainty in the preferred orientations of $s_1$ and $s_2$ is illustrated by shaded regions of azimuthal width $4\sigma$.

Finally, we assume that the encoding of the edge orientation is done through a transformation of the information contained in $\Theta$ and $\Theta - \Phi$. We model this through a function $P$ satisfying the following properties: (i) $P$ is maximized at $0°, 180°$ and is minimized at $90°$, (ii) $P$ is periodic with a period of $180°$, (iii) $P(\theta) = P(180° - \theta)$, (iv) on the interval $[0°, 90°]$, $P$ is decreasing. The first assumption models the fact that the signal is strongest when the real-world line is aligned with the preferred orientation and weakest when $90°$ apart. The second assumption is a mathematical construct allowing for the consideration of angles outside the range of $0°$ to $180°$, i.e., negative angles. The third assumption captures the observation that it is the absolute angular difference between the sensor and the edge that is encoded, not the orientated angle. The fourth assumption follows from the observation that the encoded signal weakens as the angular difference between a sensor and the edge increases.

One function that satisfies the above properties is $P(\theta) = |\cos(\theta)|$, i.e., a projection onto the horizontal axis, and thus one choice for modeling the encoded signal is given by the random variable defined by

$$\begin{aligned} S_1 &= P(\Theta) = |\cos(\Theta)|, \\ S_2 &= P(\Theta - \Phi) = |\cos(\Theta - \Phi)|. \end{aligned} \tag{4}$$

Note, since this encoding procedure will be inverted to decode the orientation of the edge, the explicit function for the encoded signal is not relevant beyond the fact that it satisfies properties (i)-(iv).

#### Decoding of Edge Orientation

The decoding problem is to use information given from signals $S_1$ and $S_2$ to determine the orientation of $E$. The challenge is that since $P(\theta) = P(180^\circ - \theta)$, the encoding function is not invertible and thus information from both signals is needed to determine the orientation of the edge. Specifically, if we treat $P^{-1}$ as a multivalued function, the decoded orientation from each signal, $\tilde{\Theta}_1 = F^{-1}(S_1)$ and $\tilde{\Theta}_2 = F^{-1}(S_2) + \Phi$, will be random variables with the following probability densities

$$\begin{aligned} \tilde{\Theta}_1 &\sim \frac{1}{2} f(\theta_1; \theta, \kappa) + \frac{1}{2} f(\theta_1; 180^\circ - \theta, \kappa), \\ \tilde{\Theta}_2 &\sim \frac{1}{2} f(\theta_2; \theta, \kappa) + \frac{1}{2} f(\theta_2; 2\phi - \theta, \kappa). \end{aligned} \tag{5}$$

Consequently, the densities of $\tilde{\Theta}_1$ and $\tilde{\Theta}_2$ are bimodal and concentrated about the correct edge orientation $\theta$ and "illusionary edges" $E_1^*$, $E_2^*$ with orientations $180^\circ - \theta$ and $2\phi - \theta$ respectively. That is, the first and second sensor would essentially decode orientations of $\theta$ or $180^\circ - \theta$ and $\theta$ or $2\phi - \theta$ with equal probability, respectively. The illusionary edges result from the fact that the reflected image of any edge about the preferred orientation of any sensor yields the same projection onto the sensor.

In Supplementary Figures 7B-7D we present a schematic diagram for the decoding of the edge orientation from signals $S_1$ and $S_2$ for varying values of $\phi$. To illustrate different possibilities we chose $\phi = 45^\circ$, $\phi = 85^\circ$, and $\phi = 3^\circ$ respectively. The light blue and pink regions illustrate the uncertainty in the decoded value of the illusionary edges, $E_1^*, E_2^*$, and each of these regions has an azimuthal width of $4\sigma$. The purple regions, overlaid on top of the light blue and pink regions, correspond to the overlap of the support of each density for the decoded random variables. Specifically, the purple region illustrates the most probable values decoded from information coming from both sensors.

To reduce the probability of decoding an illusionary edge, information from both $S_1$ and $S_2$ must be used. The difference between the individually decoded edge orientations, $u = (\tilde{\Theta}_1 - \tilde{\Theta}_2)/2$, is a natural measurement of the difference in the decoded signals. Consequently, if we assume that $u$ is close to zero, i.e., the sensors are in agreement, then $v = (\tilde{\Theta}_1 + \tilde{\Theta}_2)/2$, is a natural measurement of the fully decoded orientation. Mathematically, this decoding procedure can be made precise through computation of the joint probability density of $u$ and $v$ as well as the conditional probability that $u = 0$.

We now carry out this computation. Since we assumed $\Theta$ and $\Phi$ are independent it follows that $\tilde{\Theta}_1$ and $\tilde{\Theta}_2$ are independent. Therefore, the joint density is simply given by the product of the marginal densities:

$$(\tilde{\Theta}_1, \tilde{\Theta}_2) \sim F(\theta_1, \theta_2) = \frac{1}{4} \left( f(\theta_1; \theta, \kappa) + f(\theta_1; 180^\circ - \theta, \kappa) \right) \left( f(\theta_2; \theta, \kappa) + f(\theta_2; 2\phi - \theta, \kappa) \right). \tag{6}$$

Changing to $u, v$ coordinates, we have $d\theta_1 d\theta_2 = 2 du dv$ and thus in these coordinates the joint density is given by

$$(\tilde{\Theta}_1, \tilde{\Theta}_2) \sim G(u, v) = \frac{1}{2} \left( f(u + v; \theta, \kappa) + f(u + v; 180^\circ - \theta, \kappa) \right) \left( f(v - u; \theta, \kappa) + f(v - u; 2\phi - \theta, \kappa) \right). \tag{7}$$

Expanding and applying trigonometric identities, it follows from Equation (2) that the $u, v$ dependence can be separated as follows:

$$\begin{aligned} G(u,v) = &\frac{(180^\circ I_0(\kappa))^{-2}}{2} \left[ \exp\left( 2\kappa \cos\left( \frac{\pi(v-\theta)}{90^\circ} \right) \cos\left( \frac{\pi u}{90^\circ} \right) \right) \right. \\ &+ \exp\left( 2\kappa \cos\left( \frac{\pi(v-\phi)}{90^\circ} \right) \cos\left( \frac{\pi(u-\theta+\phi)}{90^\circ} \right) \right) \\ &+ \exp\left( 2\kappa \cos\left( \frac{\pi v}{90^\circ} \right) \cos\left( \frac{\pi(u+\theta)}{90^\circ} \right) \right) \\ &+ \left. \exp\left( 2\kappa \cos\left( \frac{\pi(v+\theta-\phi)}{90^\circ} \right) \cos\left( \frac{\pi(u+\phi)}{90^\circ} \right) \right) \right]. \end{aligned} \tag{8}$$

Consequently, the conditional probability density given $u = 0$, i.e., the sensors are in agreement, is given by

$$\begin{aligned} p(v;\theta,\phi,\kappa) =& \frac{G(0,v)}{\int_0^{180^\circ} G(0,v)dv} \\ =& C^{-1}\left(\exp\left(2\kappa\cos\left(\frac{\pi}{90^\circ}(v-\theta)\right)\right) + \exp\left(2\kappa\cos\left(\frac{\pi}{90^\circ}(v-\phi)\right)\cos\left(\frac{\pi}{90^\circ}(\theta-\phi)\right)\right)\right. \\ & \left. + \exp\left(2\kappa\cos\left(\frac{\pi}{90^\circ}v\right)\cos\left(\frac{\pi}{90^\circ}\theta\right)\right) + \exp\left(2\kappa\cos\left(\frac{\pi}{90^\circ}(v+\theta-\phi)\right)\cos\left(\frac{\pi}{90^\circ}\phi\right)\right)\right) \end{aligned}, \quad (9)$$

where the normalization constant is given by

$$C = 180^\circ\left(I_0(2\kappa) + I_0\left(2\kappa\cos\left(\frac{\pi}{90^\circ}(\theta-\phi)\right)\right) + I_0\left(2\kappa\cos\left(\frac{\pi}{90^\circ}\theta\right)\right) + I_0\left(2\kappa\cos\left(\frac{\pi}{90^\circ}\phi\right)\right)\right). \quad (10)$$

This probability density represents the distribution of decoded edge orientations from two sensors accounting for biological noise.

The probability density defined by Equation (10) is multimodal with peaks $\rho_1, \rho_2, \rho_3, \rho_4$ approximately located at

$$\begin{aligned} v_1 &= \theta, \\ v_2 &= \begin{cases} \phi & \text{if } |\theta-\phi| < 45^\circ \\ \phi + 90^\circ & \text{if } |\theta-\phi| \geq 45^\circ \end{cases}, \\ v_3 &= \begin{cases} 0^\circ & \text{if } |\theta| < 45^\circ \\ 90^\circ & \text{if } |\theta| \geq 45^\circ \end{cases}, \\ v_4 &= \begin{cases} \phi - \theta & \text{if } |\phi| < 45^\circ \\ \phi - \theta + 90^\circ & \text{if } |\phi| \geq 45^\circ \end{cases}, \end{aligned}$$

respectively. Note, the location of $\rho_1$ corresponds to the actual edge orientation while $\rho_2, \rho_3$ correspond to the orientation of the sensors, and $\rho_4$ corresponds to the orientation of the average of the two illusionary edges. Consequently, with high probability the correct orientation $\theta$ will be decoded if the relative heights $\rho_2/\rho_1$, $\rho_3/\rho_1$, and $\rho_4/\rho_1$ are all small. Approximately, we have that

$$\begin{aligned} \frac{\rho_2}{\rho_1} &\approx \begin{cases} \exp\left(2\kappa\left(1-\cos\left(\frac{\pi}{90^\circ}(\theta-\phi)\right)\right)\right) & \text{if } |\theta-\phi| < 45^\circ \\ \exp\left(2\kappa\left(1+\cos\left(\frac{\pi}{90^\circ}(\theta-\phi)\right)\right)\right) & \text{if } |\theta-\phi| \geq 45^\circ \end{cases}, \\ \frac{\rho_3}{\rho_1} &\approx \begin{cases} \exp\left(2\kappa\left(1-\cos\left(\frac{\pi}{90^\circ}\theta\right)\right)\right) & \text{if } |\theta| < 45^\circ \\ \exp\left(2\kappa\left(1+\cos\left(\frac{\pi}{90^\circ}\theta\right)\right)\right) & \text{if } |\theta| \geq 45^\circ \end{cases}, \\ \frac{\rho_4}{\rho_1} &\approx \begin{cases} \exp\left(2\kappa\left(1-\cos\left(\frac{\pi}{90^\circ}\phi\right)\right)\right) & |\phi| < 45^\circ \\ \exp\left(2\kappa\left(1+\cos\left(\frac{\pi}{90^\circ}\phi\right)\right)\right) & |\phi| \geq 45^\circ. \end{cases}. \end{aligned}$$

Therefore, it follows that $\rho_2/\rho_1$ is small unless $\theta \approx \phi$ or $\theta \approx 90^\circ + \phi$ but in either case $v_2 \approx v_1$ and thus these two peaks have merged. Similarly, $\rho_3/\rho_1$ is small unless $\theta \approx 0$ or $\theta = 90^\circ$ and thus in either case $v_3 \approx v_1$. However, if $\phi \approx 0^\circ$ or $\phi \approx 90^\circ$ we obtain $\rho_4/\rho_1 \approx 1$ and thus the illusionary edge will be decoded with a probability approximately equal to the probability of decoding the actual edge. To illustrate the behavior of the peaks of $p(v;\theta,\phi)$, in Supplementary Figure 7G we present contour plots of $p(v;\theta,\phi,\kappa)$ for various values of $\theta$. The plots validate our heuristic description in the sense that when $\phi \approx 0^\circ$ or $\phi \approx 90^\circ$ the distribution is bimodal unless $\theta \approx 0^\circ$ or $\theta \approx 90^\circ$. Moreover, when $\phi \approx \theta$ or $\phi \approx \theta + 90^\circ$ we observe that the distribution broadens out but remains centered about $\theta$. That is, the key parameter that controls the validity of the decoder is $\phi$, i.e., the angular separation between the preferred orientation of the sensors.

Motivated by the above analysis, we now develop a method for measuring decoding efficiency that is independent of the edge orientation $\theta$. Since $I_0$ grows exponentially in its argument, it follows that for values of $\theta$ bounded away from $0^\circ$ and $\phi$ that $I_0(2\kappa) \gg I_0\left(2\kappa\cos\left(\frac{\pi}{90^\circ}(\theta-\phi)\right)\right)$ and $I_0(2\kappa) \gg I_0\left(2\kappa\cos\left(\frac{\pi}{90^\circ}\theta\right)\right)$. Consequently, in this regime, the conditional density is well approximated by

$$p(v;\theta,\phi,\kappa) \approx \frac{\exp\left(2\kappa\cos\left(\frac{\pi}{90^\circ}(v-\theta)\right)\right) + \exp\left(2\kappa\cos\left(\frac{\pi}{90^\circ}(v+\theta-\phi)\right)\cos\left(\frac{\pi}{90^\circ}\phi\right)\right)}{I_0(2\kappa) + I_0\left(2\kappa\cos\left(\frac{\pi}{90^\circ}\phi\right)\right)}. \quad (11)$$

Therefore, in this regime, the probability of the decoder identifying an edge orientation that is within two standard deviations of the actual edge orientation or two standard deviations of an illusionary edge are accurately approximated by the quantities

$$\begin{aligned} p_E(\phi;\kappa) &= \int_0^{180^\circ} \frac{\exp\left(2\kappa\cos\left(\frac{\pi}{90^\circ}(v-\theta)\right)\right)}{I_0(2\kappa)+I_0\left(2\kappa\cos\left(\frac{\pi}{90^\circ}\phi\right)\right)}dv = \frac{I_0(2\kappa)}{I_0(2\kappa)+I_0\left(2\kappa\cos\left(\frac{\pi}{90^\circ}\phi\right)\right)}, \\ p_I(\phi;\kappa) &= \int_0^{180^\circ} \frac{\exp\left(2\kappa\cos\left(\frac{\pi}{90^\circ}(v+\theta-\phi)\right)\cos\left(\frac{\pi}{90^\circ}\phi\right)\right)}{I_0(2\kappa)+I_0\left(2\kappa\cos\left(\frac{\pi}{90^\circ}\phi\right)\right)}dv = \frac{I_0\left(2\kappa\cos\left(\frac{\pi}{90^\circ}\phi\right)\right)}{I_0(2\kappa)+I_0\left(2\kappa\cos\left(\frac{\pi}{90^\circ}\phi\right)\right)}. \end{aligned} \tag{12}$$

Consequently, for an arbitrary edge orientation $\theta$, since $p_E(\phi;\kappa) \geq p_I(\phi;\kappa)$ it follows that $0 \leq p_E(\phi;\kappa) - p_I(\phi;\kappa) \leq 1$ and thus the quantity $\delta = p_E(\phi;\kappa) - p_I(\phi;\kappa)$ given by

$$\delta = \frac{I_0(2\kappa) - I_0\left(2\kappa\cos\left(\frac{\pi}{90^\circ}\phi\right)\right)}{I_0(2\kappa)+I_0\left(2\kappa\cos\left(\frac{\pi}{90^\circ}\phi\right)\right)} \tag{13}$$

is a measurement of the difference in probability of decoding the correct edge orientation vs decoding the illusionary edge. We call $\delta$ the decoding efficiency. In Supplementary Figure 7H we present a plot of the decoding efficiency versus the angular separation between the sensors. For the level of biological noise we considered, this plot indicates that this decoding procedure will robustly recover approximately the correct edge orientation for values of $\phi$ between $10^\circ$ and $80^\circ$. In Supplementary Figure 7I we illustrate the how this efficiency changes with different levels of biological noise.
